# Allele-specific expression reveals complex *cis*- and *trans*-regulatory divergence underlying fruit phenotypic differences between cultivated and wild tomato species

**DOI:** 10.64898/2026.05.27.728279

**Authors:** Jiantao Zhao, Philippe Nicolas, Yimin Xu, Julia Vrebalov, James J. Giovannoni, Zhangjun Fei, Carmen Catala

**Author notes:** Present address: Key Laboratory of Genome Research and Genetic Improvement of Xinjiang Characteristic Fruits and Vegetables, Institute of Fruits and Vegetables, Academy of Agricultural Sciences of Xinjiang Uygur Autonomous Region, Urumqi, China (Jiantao Zhao). Present address: Department of Plant and Soil Science, Texas Tech University, Lubbock, TX, USA. These authors contributed equally (Jiantao Zhao, Philippe Nicolas).

## Abstract

Tomato (*Solanum lycopersicum*) is one of the most important agricultural crops and serves as a model system for fleshy fruit biology. However, domestication bottlenecks have led to limited genetic diversity for further crop improvement. Wild relatives represent a rich reservoir of phenotypic diversity, yet the molecular basis underlying such phenotypic diversity remains largely unknown. As gene expression variation is a major driver of phenotypic diversity, we performed genome-wide allele-specific expression analyses in F_1_ hybrids between cultivated tomato and three wild relatives spanning a range of evolutionary distances, across three fruit tissues and multiple developmental stages. Our analysis generated a multi-species map of *cis* – and *trans*-regulatory variation, revealing a predominant role of *cis*-regulatory effects in expression divergence across species, tissues, and stages. The majority of *cis* effects were tissue– and/or stage-specific, underscoring the importance of tissue context in regulatory variation. Regulatory mechanisms and inheritance patterns shifted with evolutionary distance, with more distantly related species showing increased *cis*-regulatory contributions. Finally, we identified extensive regulatory divergence in biosynthetic pathways related to fruit nutritional value and flavor quality, including carotenoids, flavonoids, alkaloids, sugars, and volatiles. This study presents a high-resolution map of regulatory variation underlying tomato fruit development and provides evolutionary insights into the regulation of fruit nutrition and flavor traits.

## Introduction

Cultivated tomato (*Solanum lycopersicum*), one of the most important agricultural crops worldwide, has undergone domestication and intensive breeding to improve traits such as yield, fruit size, color, firmness, and plant architecture, resulting in high-performing modern varieties (Bergougnoux, 2014; Lin et al., 2014). However, domestication bottlenecks have caused a substantial loss of genetic diversity, limiting the potential for further improvement of cultivated germplasm (Bai and Lindhout, 2007; Razifard et al., 2020). In contrast, wild tomato species display remarkable morphological and metabolic diversity and represent a rich reservoir of genetic variation that can be harnessed to improve agronomic traits such as nutritional and flavor quality, yield, and tolerance to biotic and abiotic stresses (Soyk et al., 2017; Tieman et al., 2017; Zhao et al., 2019; Li et al., 2023). Recent studies employing transcriptomics, metabolomics, genome-wide association studies (GWAS), and pangenomics have uncovered multiple loci and structural variants underlying fruit trait diversity (Zhu et al., 2018; Zhao et al., 2019; Alonge et al., 2020; Szymański et al., 2020; Wang et al., 2020; Li et al., 2023). Despite these advances, the molecular basis underlying phenotypic differences between cultivated and wild tomato species remains largely unknown.

Phenotypic differences between species can arise from genetic variation in coding sequences as well as from differences in gene expression. Increasing evidence suggests that changes in gene expression contribute substantially to phenotypic evolution and speciation (Tiroshauth et al., 2009; Meyer and Purugganan, 2013; Lemmon et al., 2014; Alonge et al., 2020). In tomato, several genes cause phenotypic differences as a result of changes in gene regulation rather than protein function. Examples include genes controlling fruit size and shape (Cong et al., 2002; Xiao et al., 2008; Chakrabarti et al., 2013), flowering induction (Soyk et al., 2017), style length (Chen et al., 2007), and flavor-related volatiles (Gao et al., 2019).

Regulatory divergence leading to gene expression differences can be caused by *cis*-acting factors (e.g., DNA polymorphisms or DNA methylation changes), *trans*-acting factors (e.g., transcription factors or regulatory RNAs), or a combination of *trans-* and *cis*-effects (Wittkopp et al., 2004; Tiroshauth et al., 2009; McManus et al., 2010; Lemmon et al., 2014; Crowley et al., 2015; Signor and Nuzhdin, 2018; Bao et al., 2019). Mutations in *cis*-regulatory sequences represent a prevalent mechanism underlying the evolution of phenotypic diversity, driving natural or domestication-associated selection (Wittkopp and Kalay, 2012; Meyer and Purugganan, 2013; Lemmon et al., 2014). Compared with mutations in coding regions, changes in *cis*-regulatory elements are often less pleiotropic and can affect not only gene expression levels but also the spatial and temporal specificity of expression. Consequently, *cis*-regulatory variants represent attractive targets for crop improvement (Swinnen et al., 2016; Rodríguez-Leal et al., 2017).

Comparative transcriptomic analyses have revealed extensive gene expression differences associated with phenotypic variation between cultivated tomato and its wild relatives (Koenig et al., 2013; Sauvage et al., 2017; Doron-Faigenboim et al., 2023). However, such studies offer limited insight into the regulatory mechanisms underlying gene expression divergence. One approach to differentiate between *cis*– and *trans*-regulatory effects is the analysis of allele-specific expression (ASE) in F_1_ hybrids compared with their parents. In F_1_ hybrids, the two parental alleles are exposed to the same cellular environment and therefore to the same set of *trans*-acting factors. Consequently, ASE differences in F_1_ hybrids provide a readout of *cis*-regulatory activities, whereas expression differences between the parents that are not observed in the hybrids indicate *trans*-regulatory divergence (Signor and Nuzhdin, 2018). Genome-wide ASE analyses have been used to characterize *cis-* and *trans*-regulatory divergence in several plant species (Cubillos et al., 2014; Lemmon et al., 2014; Shao et al., 2019; Shi et al., 2024). In tomato, ASE analysis has been used to characterize gene expression differences between cherry-type and large-fruited accessions (Albert et al., 2018), and it has yet to be applied to evaluate interspecific gene expression variation between cultivated tomato and its wild relatives.

In recent years, quantitative trait locus (QTL) mapping, GWAS, and pangenome studies have uncovered that many genetic changes associated with tomato domestication and breeding reside in *cis*-regulatory regions (Gao et al., 2019; Alonge et al., 2020; Szymański et al., 2020; Wang et al., 2020; Li et al., 2023). Nevertheless, the number of *cis*-regulated alleles identified to date remains limited. This underscores the need for systematic approaches to expand the identification and characterization of regulatory variants in tomato‒not only to elucidate the genetic basis of phenotypic divergence between cultivated tomato and its wild relatives but also to support improvement of fruit traits in cultivated tomato.

In this study, we performed a tissue-specific ASE analysis across tomato fruit development using F_1_ hybrids between cultivated tomato and three wild relatives spanning different evolutionary distances. To enhance the power of detecting regulatory variants, we analyzed ASE in three fruit tissues collected at three or four developmental stages. We examined both *cis-* and *trans*-regulatory effects, their interactions, and patterns of gene expression inheritance. Our results reveal a significant role of *cis*-regulatory divergence in determining gene expression differences between cultivated and wild tomato species. The majority of *cis* effects were specific to particular fruit tissues and/or developmental stages, underscoring the importance of tissue context in regulatory variation. Regulatory variation was identified in diverse metabolic pathways, providing evolutionary insights into the regulation of fruit nutritional quality and flavor traits. This study provides a comprehensive inventory of allele-specific gene expression that can be leveraged to explore regulatory divergence in genes and pathways contributing to fruit phenotypic diversity.

## Results

### Transcriptome sequencing of F_1_ hybrids and genome assemblies of parental lines

To investigate the basis of gene expression divergence in tomato, we performed ASE analysis using F_1_ hybrids between *S. lycopersicum* and three wild tomato species that represent distinct evolutionary lineages: the red-fruited close relative *S. pimpinellifolium* and the more distantly related green-fruited species *S. neorickii* and *S. pennellii*. Crosses were made between each wild species and its respective *S. lycopersicum* line‒NC EBR-1 × *S. pimpinellifolium* LA2093, TA209 × *S. neorickii* LA2133, and M82 × *S. pennellii* LA0716‒with reciprocal hybrids generated for the NC EBR-1 × *S. pimpinellifolium* LA2093 cross (**Fig. 1a**). Parental lines were selected to align with introgression lines (ILs) for *S. pennellii* (Alseekh et al., 2013), recombinant inbred lines (RILs) for *S. pimpinellifolium* (Ashrafi et al., 2012), and backcross inbred lines (BILs) for *S. neorickii* (Brog et al., 2019). RNA-seq analysis was performed for three fruit tissues (pericarp, placenta, and jelly) at multiple developmental stages spanning fruit growth and maturation in both parents and F_1_ hybrids, with three biological replicates per sample. On average, 39.5 million and 19.0 million cleaned read pairs were generated for the F_1_ hybrids and the parents, respectively (**Supplementary Table 1**).

**Fig. 1.**
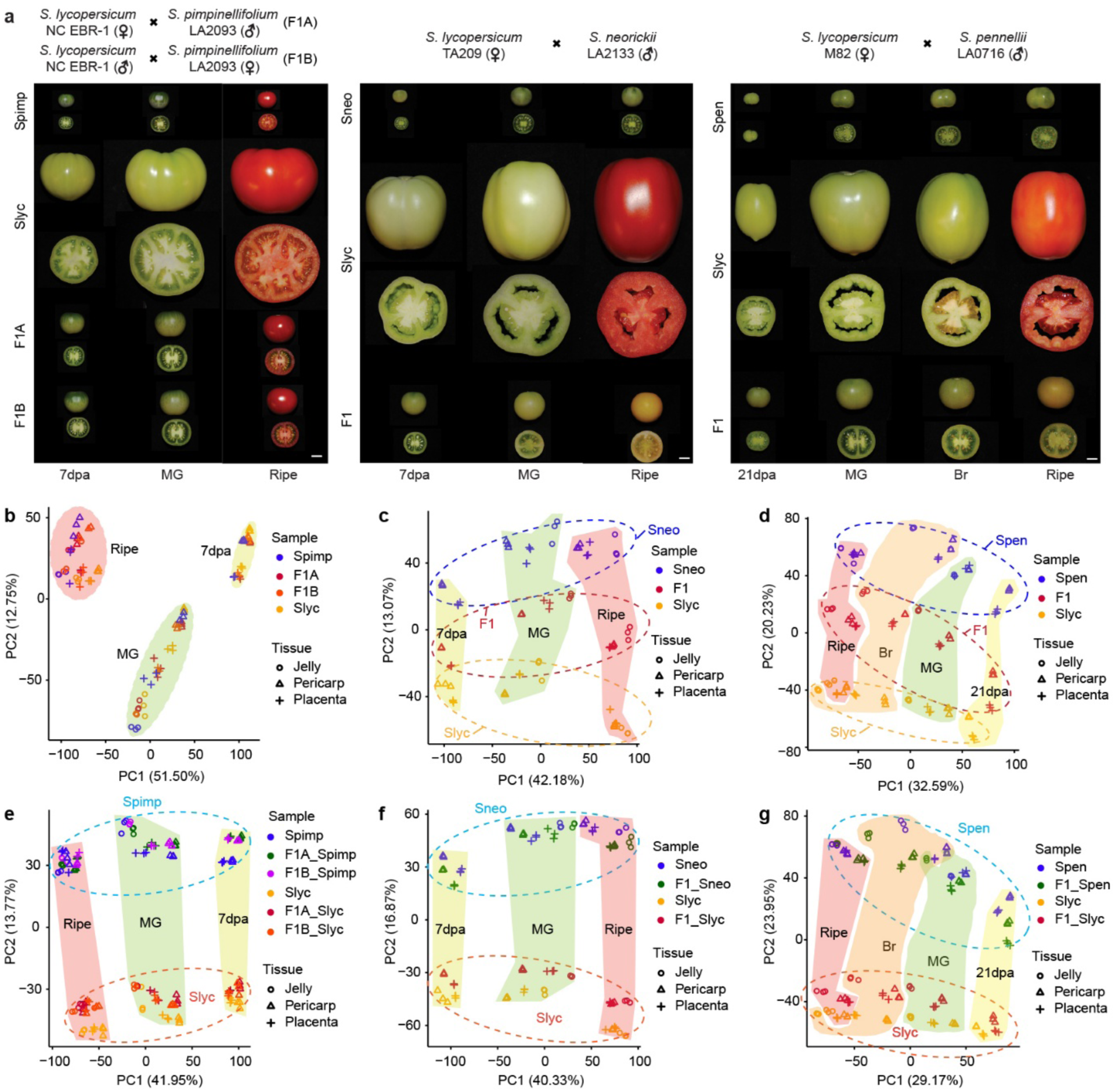
Experimental design and principal component analysis (PCA) of gene expression across genotypes, fruit tissues, and developmental stages. **a**, Schematic of crosses and representative fruit images of the parental lines and corresponding F_1_ hybrids. b-d, PCA of total read counts for parents and F_1_ hybrids derived from crosses between *S. lycopersicum* and *S. pimpinellifolium* (b), *S. neorickii* (c), and *S. pennellii* (d). e-g, PCA of allele-specific read counts for parents and F_1_ hybrids from crosses between Slyc and Spimp (e), Sneo (f), and Spen (g). dpa, days post anthesis; MG, mature green; Br, breaker. Slyc, *S. lycopersicum*; Spimp, *S. pimpinellifolium*; Sneo, *S. neorickii*; Spen, *S. pennellii*. Scale bar = 1 cm.

To enable accurate ASE quantification, we leveraged high-quality genome assemblies and annotations for all parental lines. Reference genomes were already available for *S. pimpinellifolium* LA2093 (Wang et al., 2020) and *S. lycopersicum* M82 (Zhou et al., 2022). For *S. neorickii* LA2133 and *S. pennellii* LA0716, we generated *de novo* assemblies using PacBio high-fidelity (HiFi) long reads. We obtained 32.72 Gb of HiFi data (34.4× coverage, mean read length of 13.2 kb) for LA2133 and 22.85 Gb (19.0× coverage, mean read length of 13.4 kb) for LA0716. The resulting assemblies were highly contiguous, with total sizes of 831.2 Mb (contig N50 = 8.3 Mb) for LA2133 and 1,062.2 Mb (contig N50 = 10.6 Mb) for LA0716, and with 97.45% and 95.51% of contigs anchored to chromosomes, respectively (**Supplementary Table 2**). High genome completeness and accuracy were confirmed by k-mer analysis using Merqury (Rhie et al., 2020) and BUSCO assessments (Simão et al., 2015) (**Supplementary Tables 2** and **3**). For the *S. lycopersicum* parental accessions NC EBR-1 and TA209, we reconstructed pseudogenomes using Illumina short-read sequencing (64.4 Gb, 67.8× for NC EBR-1; 57.9 Gb, 60.9× for TA209), aligning reads to the ‘Heinz 1706’ reference genome (v4.0) (Hosmani et al., 2019).

Synteny-guided orthologue mapping identified 37,686, 34,826, and 38,611 one-to-one gene pairs between the parental genomes of *S. lycopersicum* and *S. pimpinellifolium*, *S. lycopersicum* and *S. neorickii*, and *S. lycopersicum* and *S. pennellii*, respectively (**Supplementary Table 4**). As expected, the number of parent-specific genes increased with increasing phylogenetic distance from *S. lycopersicum*.

### Gene expression patterns in F_1_ hybrids and their parental species

Principal component analysis (PCA) of genome-wide expression profiles revealed that *S. lycopersicum* and *S. pimpinellifolium* clustered closely with their F_1_ hybrids, with developmental stage representing the major variable separating samples. Notably, at the ripe stage, pericarp samples from the hybrids shifted toward the *S. pimpinellifolium* parent, indicating subtle parental influences on ripening-related transcriptional programs (**Fig. 1b**). In contrast, hybrids involving the more distantly related *S. neorickii* and *S. pennellii* separated clearly from their respective parental lines, in addition to the separation by developmental stage (**Fig. 1c,d**). Tissue-specific divergence was also observed; for example, jelly samples at the mature green (MG) stage in *S. neorickii* hybrids and parents, and jelly sample at the breaker (Br) stage in *S. pennellii* hybrids and parents, were clearly separated from the other samples (**Fig. 1c,d**).

To quantify allele-specific expression across different fruit tissues and developmental stages, RNA-seq reads from F_1_ hybrids and their parents were partitioned into parent-specific categories based on alignments to their respective parental genomes (**Supplementary Table 1**). PCA of allele-specific expression profiles revealed consistent separation between parental alleles, with F_1_ hybrid alleles clustering alongside their corresponding parents (**Fig. 1e-g**). Developmental stage emerged as the primary factor shaping allele-specific expression in F_1_ hybrids and parents, outweighing tissue-specific effects. However, notable exceptions were observed. For instance, in *S. lycopersicum* × *S. pennellii* hybrids, the *S. pennellii* allele in jelly tissue showed partial separation from other tissues at the MG and Br stages, highlighting localized regulatory divergence restricted to specific tissue–stage combinations (**Fig. 1g**). Sample correlation analysis of allele-specific gene expression further confirmed that parental alleles and their corresponding alleles in F_1_ hybrids were in general tightly clustered according to developmental stage (**Supplementary Fig. 1**). Within stages, tissues generally clustered together, though discrepancies were observed at the ripe stage, where jelly, pericarp, and placenta occasionally co-clustered, reflecting partial convergence of transcriptional programs during ripening. Collectively, these results demonstrate that developmental stage is the primary determinant of transcriptional regulation in tomato hybrids while tissue type exerts a secondary yet still meaningful influence, particularly during early ripening stages.

### Regulatory effects underlying divergence between cultivated and wild tomato species

We next explored transcriptional divergence between cultivated tomato and its wild relatives by assigning differentially expressed genes to different regulatory categories following established frameworks (McManus et al., 2010; Lemmon et al., 2014), focusing on *cis*, *trans*, and their interactions (*cis* + *trans* or *cis* × *trans*). Approximately half of the annotated genes were expressed in each tissue and developmental stage in all hybrids (**Supplementary Table 5**). The majority exhibited conserved expression (no significant allelic expression differences between parents or in hybrids), with the highest proportions observed in *S. pimpinellifolium* hybrids (55-75%), followed by *S. neorickii* (52-61%) and *S. pennellii* hybrids (42-56%) (**Supplementary Table 5**). In the reciprocal *S. lycopersicum* × *S. pimpinellifolium* hybrids, 6,353 and 6,677 genes showed evidence of *cis* and/or *trans* effects **(Supplementary Data 1** and **2**), whereas nearly twice as many genes exhibited such effects in *S. neorickii* (12,716) and *S. pennellii* (14,460) hybrids (**Supplementary Data 3 and 4)**. This pattern indicates that regulatory divergence increases with increasing evolutionary distance from cultivated tomato. Reciprocal *S. lycopersicum* × *S. pimpinellifolium* hybrids exhibited nearly identical proportions of genes across regulatory categories, indicating negligible parent-of-origin effects (**Fig. 2; Supplementary Fig. 2**).

**Fig. 2.**
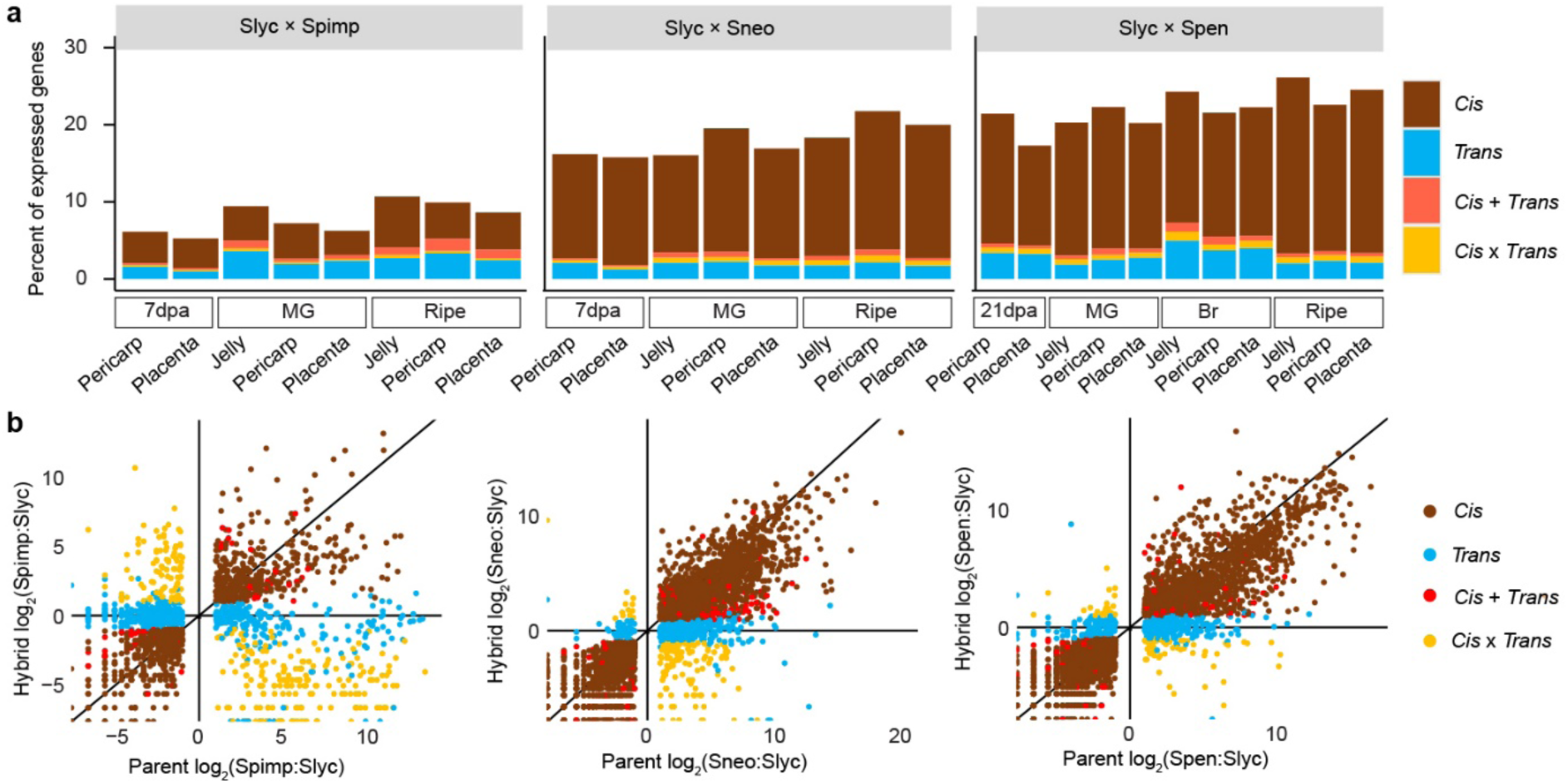
Allele-specific expression in tomato hybrids. **a**, Percentage of genes in different regulatory categories across fruit tissues and developmental stages. **b**, Scatter plots showing the relative expression of cultivated and wild alleles in parents versus the relative expression of the corresponding alleles in the F_1_ hybrid for pericarp tissue at the ripe stage. MG, mature green; Br, breaker. Slyc, *S. lycopersicum*; Spimp, *S. pimpinellifolium*; Sneo, *S. neorickii*; Spen, *S. pennellii*.

In all tissues and stages *cis*-regulated genes were prevalent among those showing allelic expression differences, accounting for 3-22% of expressed genes, with notably higher proportions in *S. neorickii* and *S. pennellii* hybrids (**Fig. 2a,b**). Overall, the number of *cis*-regulated genes increased during ripening (**Fig. 2a**; **Supplementary Table 5**), suggesting a central role of *cis*-acting variants in fine-tuning ripening-associated pathways. In contrast, *trans*-regulated genes represented a smaller proportion (1-5% of expressed genes) and displayed a relatively consistent pattern across hybrids. Genes with combined *cis-* and *trans*-regulation (0.4-2%) or compensatory effects (0.4-1.8%) were comparatively rare. Tissue-level analyses revealed slightly higher proportions of *cis*-regulated genes in jelly compared to placenta and pericarp in ripe tissues of *S. pimpinellifolium* and *S. pennellii* hybrids, whereas in *S. neorickii* hybrids, pericarp harbored a higher proportion of *cis*-regulated genes than jelly and placenta (**Fig. 2a**; **Supplementary Table 5**).

To determine the relative contributions of *cis* and *trans* regulation to parental expression divergence, we calculated the proportion of divergence due to *cis* effects for groups of genes binned by the magnitude of parental expression differences calculated as |log_2_ (P2/P1)| (P1, cultivated parent; P2, wild parent). In general, divergence due to *cis* effects increased with total parental divergence in all hybrids, with the exception of *S.pimpinellifolium* hybrids at the ripe stage (**Fig. 3a**; **Supplementary Fig. 3a**). The increase in the proportion of *cis* regulatory contributions with greater parental divergence was particularly pronounced for genes under *cis* + *trans* regulation, in which *cis* and *trans* effects acted in the same direction (**Supplementary Fig. 4**). Quantification of parental expression divergence magnitude, further showed that *cis* + *trans*-regulated genes consistently exhibited the largest parental expression differences in both *S. neorickii* and *S. pennellii* hybrids (**Fig. 3b**; **Supplementary Fig. 3b**). In *S. pimpinellifolium* hybrids, the largest parental expression differences were generally observed in either *cis*, *cis* + *trans* or *cis* × *trans*-regulated genes, depending on tissue and developmental stage (**Fig. 3b**; **Supplementary Fig. 3b**). A striking phylogenetic pattern emerged in parental expression bias expressed as log_2_(P2/P1). Both *S. neorickii* and *S. pennellii* hybrids displayed predominant upregulation of wild alleles (parental divergence direction > 0) across all regulatory modes, whereas in *S. pimpinellifolium* hybrids this bias was restricted to specific tissues and developmental stages (**Fig. 3c**; **Supplementary Fig. 3c**). As expected, genes with *cis*-regulated effects (*cis*, *cis* + *trans*, or *cis* × *trans*) exhibited greater *cis*-regulatory divergence magnitude than *trans* effect in F_1_ hybrids, which showed near zero *cis* effect magnitude (**Fig. 3d**; **Supplementary Fig. 3d**). Further examination of *cis*-regulatory divergence revealed consistent directional biases: genes under *cis* and *cis* + *trans* regulation preferentially showed upregulation of the wild allele (*cis* divergence direction > 0), albeit with minor discrepancies in specific tissues or stages (**Fig. 3e**; **Supplementary Fig. 3e**). In contrast, *cis* × *trans*-regulated genes exhibited predominantly upregulation of cultivated alleles (*cis* divergence direction < 0) across most tissues and hybrids, with exceptions in tissues at 7 dpa and MG in *S. pimpinellifolium* hybrids (**Fig. 3e**; **Supplementary Fig. 3e**). Collectively, these findings demonstrate that *cis*-regulatory contributions increase with overall parental expression divergence, with *cis* + *trans* regulation driving the most pronounced transcriptional differences and developmental stage dynamically modulating parental expression biases.

**Fig. 3.**
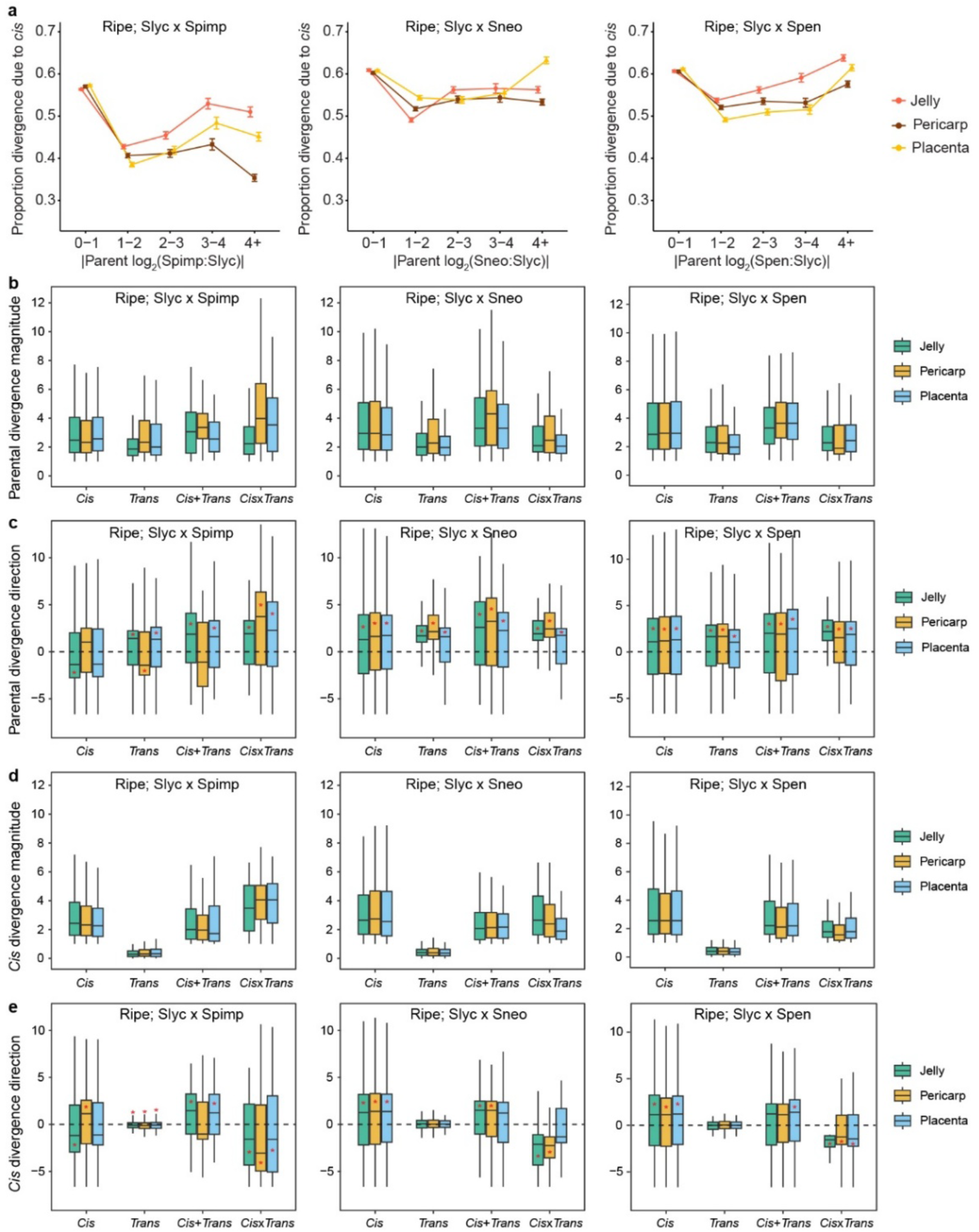
Expression divergence attributable to *cis* regulation at the ripe stage. **a**, Proportion of parental expression divergence explained by *cis*-regulatory effects. Genes were binned by absolute parental divergence in x-axis. **b**-**e**, Parental divergence magnitude [|log_2_(P2/P1)|] (**b**), parental divergence direction [log_2_(P2/P1)] (**c**), *cis* divergence magnitude [|log_2_(P2/P1)| in F_1_] (**d**), and *cis* divergence direction [log_2_(P2/P1) in F] (**e**) for genes under *cis*, *trans*, *cis* + *trans*, or *cis* × *trans* regulation. P1, cultivated parent; P2, wild parent. In all panels: *S. lycopersicum* × *S. pimpinellifolium* (left), *S. lycopersicum* × *S. neorickii* (middle) and *S. lycopersicum* × *S. pennellii* (right). Significant deviations from zero, indicated by red stars (*), were inferred using the Wilcoxon test (*P* < 0.05).

An analysis of the tissue– and stage-specific distribution of *cis*-regulated genes revealed that the proportion of genes exhibiting *cis* effects exclusively in a single tissue and developmental stage ranged from 3% to 38% (**Supplementary Table 6**). The proportion of tissue-specific *cis*-regulated genes decreased with increasing phylogenetic distance from cultivated tomato and was overall lower in the placenta than in other tissues (**Fig. 4**). Although most tissue-specific *cis*-regulated genes were unique to individual hybrids, some showed conserved tissue-specific *cis* effects across all hybrids. For example, *SlKLUH* (*Solyc03g114940*), which encodes a cytochrome P450 involved in controlling fruit mass (Chakrabarti et al., 2013), displayed placenta-specific *cis* regulation in all hybrids, and was expressed predominantly at early developmental stages. Expression of the wild allele of *SlKLUH* was significantly lower than that of the *S. lycopersicum* allele (**Supplementary Fig. 5a**), consistent with the reported association between reduced *SlKLUH* expression and smaller fruit size (Chakrabarti et al., 2013; Alonge et al., 2020). Another example is a zeta-carotene desaturase gene (*Solyc01g097810*), which in *S. pennellii* hybrids exhibited *cis* regulation exclusively in the pericarp at the ripe stage, despite being expressed in all fruit tissues (**Supplementary Fig. 5b)**. Notably, pericarp-specific *cis* regulation of this gene was also observed in *S. pimpinellifolium* hybrids; however, in this case expression was biased toward the wild allele, whereas in *S. pennellii* hybrids the wild allele was downregulated (**Supplementary Fig. 5b**). Together these examples underscore the remarkable tissue specificity of *cis*-regulatory events.

**Fig. 4.**
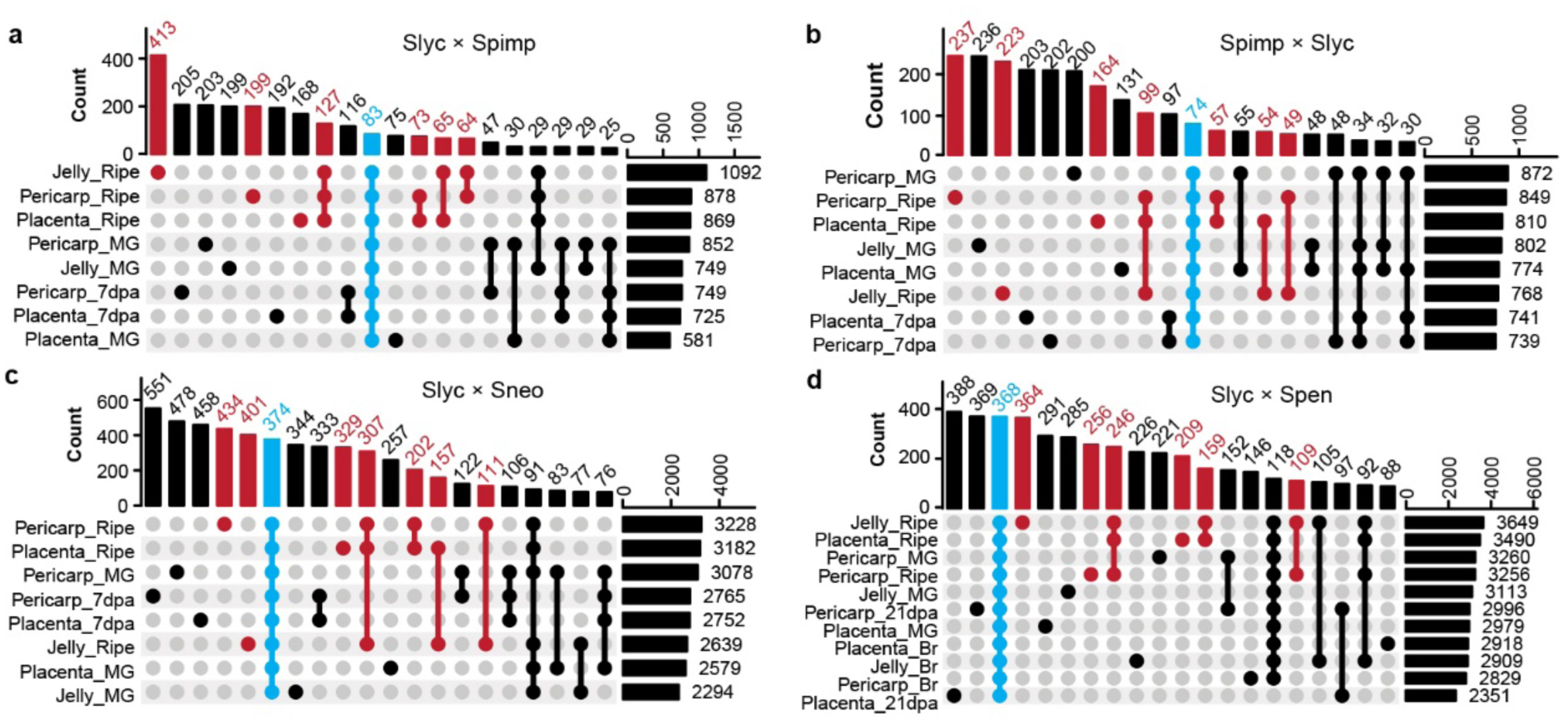
Tissue– and stage-specific distribution of *cis*-regulated genes. **a-d**, Upset plot of *cis*-regulated genes from *S. lycopersicum* × *S. pimpinellifolium* (**a**), *S. pimpinelifolium* × *S. lycopersicum* (**b**), *S. lycopersicum* × *S. neorickii* (**c**), and *S. lycopersicum* × *S. penellii* (**d**). *Cis*-regulated genes at the ripe stage are in dark red. Genes showing *cis*-regulation in all tissues and stages are in cyan. dpa, days post anthesis; MG, mature green; Br, breaker.

To explore the genetic basis of these *cis* effects, we examined the enrichment of structural variants (SVs; ≥30 bp) within 2-kb promoter regions of *cis-* versus *trans-*regulated genes. In *S. pimpinellifolium* hybrids, *cis*-regulated genes were more likely to harbor promoter SVs than *trans*-regulated genes, whereas in *S. neorickii* and *S. pennellii* hybrids, the proportions were more balanced (**Supplementary Table 7**). Notably, among constitutively *cis*-regulated genes‒those showing *cis* effects across all tissues and stages‒promoter SVs were strikingly enriched, particularly in *S. pennellii* hybrids, where 70.7% of such genes carried them (**Supplementary Table 7**). This suggests that promoter SVs are likely a key driver of stable *cis*-regulatory divergence, especially in the more distantly related wild species.

### Gene expression inheritance in F_1_ hybrids

We next examined gene expression inheritance patterns in the hybrids and classified them as additive, dominant, overdominant, and underdominant. Across all hybrids, the total numbers of dominant and overdominant genes were similar, but notable differences were observed in the numbers of underdominant and additive genes (**Supplementary Fig. 6**). Specifically, *S. neorickii* and *S. pennellii* hybrids contained fewer underdominant genes and showed a substantially higher prevalence of additive inheritance compared with *S. pimpinellifolium* hybrids. Tissue– and stage-specific patterns were also evident. For instance, jelly samples at the MG stage in *S. pimpinellifolium* hybrids and placenta samples at 21 dpa in *S. pennellii* hybrids, harbored higher proportion of cultivated parent dominant genes (P1-dominant) relative to other tissues. More broadly, wild parent dominance (P2-dominant) was predominant in *S. neorickii* and *S. pennellii* hybrids (**Fig. 5a**), suggesting that with increasing evolutionary divergence, transcriptional control in hybrids becomes increasingly biased toward wild alleles.

**Fig. 5.**
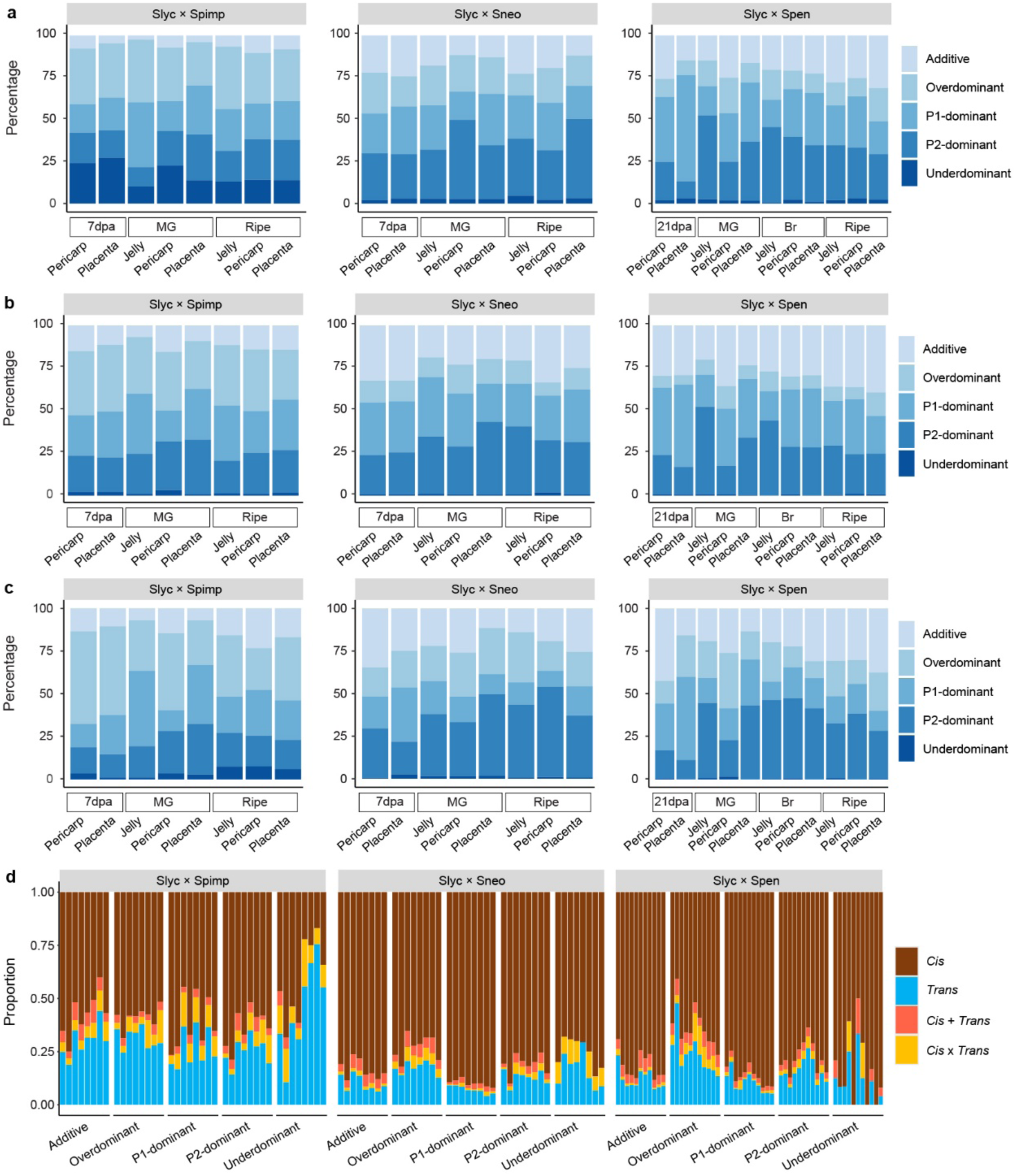
Gene expression inheritance in tomato hybrids. **a**, Percentage of genes in each inheritance category across fruit tissues and developmental stages. **b,c**, Distribution of inheritance modes among *cis-*(**b**) and *trans*-regulated genes (**c**). **d**, Proportion of genes showing *cis*, *trans, cis + trans*, and *cis × trans* regulation for each inheritance category. Within each inheritance category, bars represent individual stages or tissues, shown in the same order as in panel **a**. P1, cultivated parent; P2, wild parent. dpa, days post anthesis; MG, mature green; Br, breaker. Slyc, *S. lycopersicum*; Spimp, *S. pimpinellifolium*; Sneo, *S. neorickii*; Spen, *S. pennellii*.

To establish functional connections between inheritance pattern and regulatory mechanisms, we systematically examined distributions of inheritance modes among *cis*– and trans-regulated genes. In *S. pimpinellifolium* hybrids, *cis*– regulated genes predominantly exhibited over-dominant inheritance (28.2-39.2%, average 33.7%) and markedly low additive inheritance (6.8-15.5%, average 12.3%) (**Fig. 5b; Supplementary Fig. 6**). This regulatory profile shifted in hybrids with more phylogenetically distant species: *S. neorickii* hybrids showed a marked increase in additive inheritance (18.8-33.5%, average 25.7%) accompanied by a reduction in over-dominance (average 12.8%). Notably, *S. pennellii* hybrids displayed extreme parental-dominance biases, with P1-dominant inheritance reaching 48.3% in placenta tissue at 21 dpa, and P2-dominant inheritance peaking at 51.4% in jelly tissue at the MG stage, while intermediate additive inheritance levels were maintained (20.1-35.7%, average 27.9%) (**Fig. 5b; Supplementary Fig. 6**).

Inheritance patterns for *trans*-regulated genes in *S. pimpinellifolium* hybrids were similar to those observed for *cis*-regulated genes and showed pronounced over-dominance (24.38-54.06%; mean: 37.95%), exceeding the levels in *cis*-genes by ∼4%. Both *S. neorickii* and *S. pennellii* hybrids exhibited elevated P2-dominant inheritance compared to *S. pimpinellifolium* (**Fig. 5c**). Collectively, these data reveal an inverse relationship between phylogenetic distance and over-dominant inheritance, with increasing parental-dominance polarization in more distant relatives, suggesting that transcriptional inheritance architectures are systematically reprogrammed with genetic divergence.

We further evaluated the contribution of *cis* regulation across inheritance categories. In *S. neorickii* and *S. pennellii* hybrids, *cis*-regulated genes were prevalent across all inheritance categories (**Fig. 5d**), consistent with the pervasive role of *cis* variation in shaping gene expression divergence in these lineages (**Fig. 2a**). By contrast, in *S. pimpinellifolium* hybrid*s*, *cis-* and *trans*-regulation contributed more evenly across categories, with one exception: underdominant inheritance at the ripe stage showed an increased proportion of *trans-*regulated genes. Altogether, these results indicate that, with increasing phylogenetic distance, gene expression inheritance in tomato hybrids shifts towards additive and wild-parent dominant modes, driven largely by *cis*-regulatory changes, providing mechanistic insight into the regulatory complexity underlying phenotypic diversity.

### Regulatory variation underlying tomato flavor and nutrition

To investigate how regulatory divergence shapes tomato fruit nutritional and flavor quality, we examined allele-specific expression of genes involved in the accumulation of carotenoids, flavonoids, alkaloids, sugars, and volatiles—key determinants of tomato nutrition, flavor, and consumer preference.

Carotenoids provide both pigmentation and nutritional values. Unlike red-fruited species, *S. pennellii* and *S. neorickii* retain green fruit at the ripe stage due to the lack of lycopene accumulation. Key genes—including *GGPPS* (*Solyc04g079960*, *Solyc09g008920*), *PSY1* (*Solyc03g031860*), *PDS* (*Solyc03g123760*), *Z-ISO* (*Solyc12g098710*), *ZDS* (*Solyc01g097810*), *LCYB* (*Solyc10g079480*), and *LCYE* _(_*Solyc12g008980*)—displayed lineage-specific regulation dominated by *cis*-effects (**Fig. 6a**). Hybrids with the green-fruited species showed *cis* and *trans* effects upregulating the cultivated alleles of *PSY1* and genes involved in downstream steps of lycopene synthesis, such as *Z-ISO*. In contrast, the wild alleles of *LCYE* and *LCYB*, which divert carotenoid flux away from lycopene toward δ– and γ-carotenes, were upregulated exclusively through *cis* effects (**Supplementary Fig. 7**). This allele-specific rewiring of the carotenoid pathway explains the lack of lycopene accumulation in green-fruited wild species and illustrates how *cis*-regulatory changes drive striking phenotypic divergence in fruit color and nutritional quality.

**Fig. 6.**
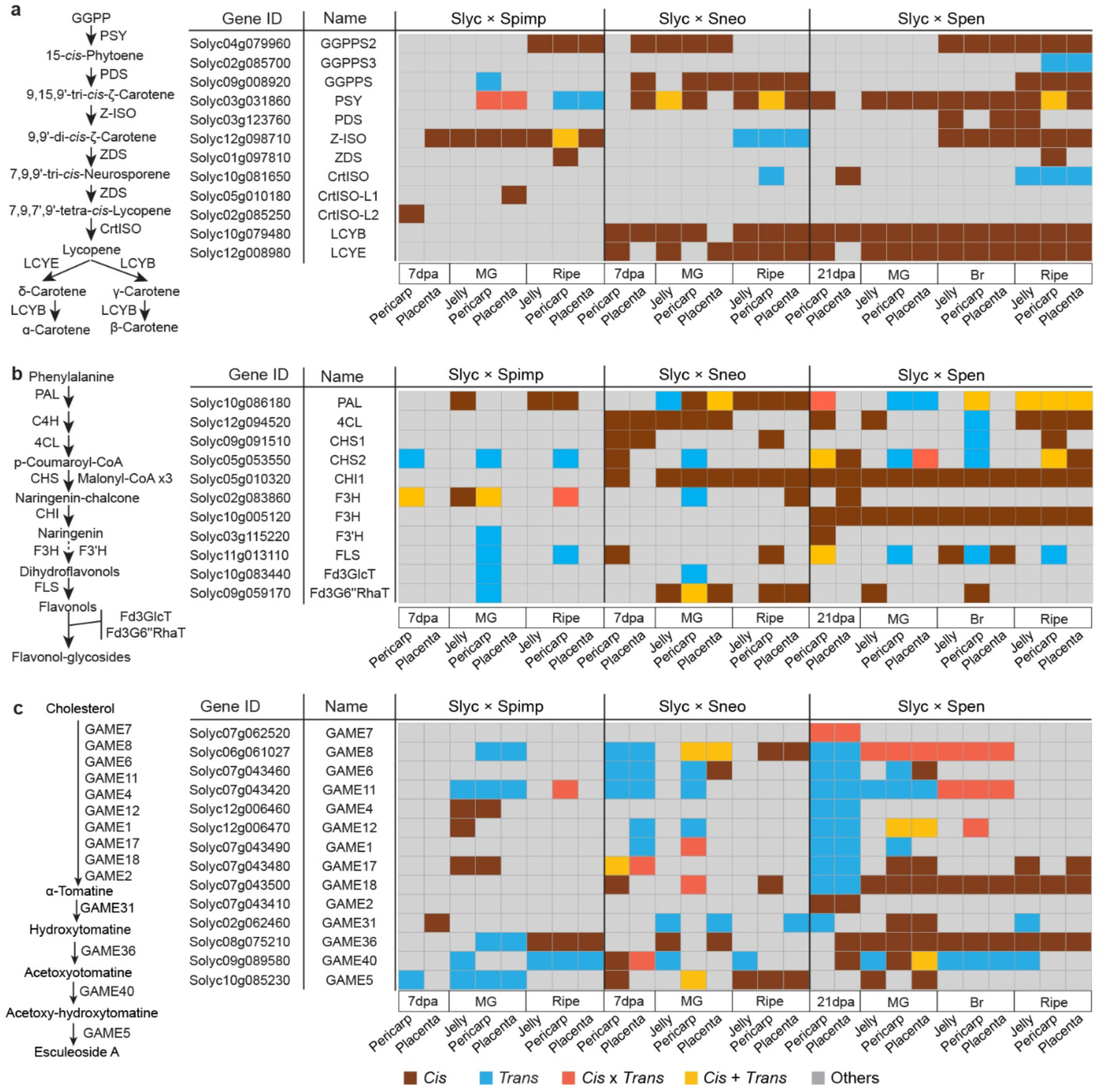
Regulatory patterns of nutritional and flavor traits in tomato hybrids. **a**, Pathway and regulatory patterns of carotenoid biosynthesis. **b**, Pathway and regulatory patterns of phenylpropanoid and flavonoid biosynthesis. **c**, Pathway and regulatory patterns of steroidal glycoalkaloid biosynthesis. The “Others” regulatory category included conserved, ambiguous, compensatory, and non-expressed. GGPPS, geranylgeranyl diphosphate synthase; PSY1, phytoene synthase 1; PDS, phytoene desaturase; Z-ISO, ζ-carotene isomerase; ZDS, ζ-carotene desaturase; CRTISO, carotenoid isomerase; LCYE, lycopene ε-cyclase; LCYB, lycopene β-cyclase; PAL, phenylalanine ammonia-lyase; 4CL, 4-coumarate: CoA ligase; CHS, chalcone synthase; CHI, chalcone isomerase; F3H, flavanone 3-hydroxylase; F3’H, flavonoid 3’-hydroxylase; FLS, flavonol synthase; Fd3GlcT, flavonoid 3-O-glucosyltransferase; Fd3G6’’RhaT, flavonoid 3-O-glucoside, 6’’-O-rhamnosyltransferase; GAME, glycoalkaloid metabolism.

We observed widespread *cis* or *cis + trans* regulation of phenylpropanoid pathway genes in *S. neorickii* and *S. pennellii* hybrids (**Fig. 6b**). For example, *PAL* (*Solyc10g086180*), encoding phenylalanine ammonia lyase, a key enzyme regulating hydroxycinnamate and flavonoid accumulation (Brog et al., 2019), showed near-constitutive *cis*-regulation with significantly higher expression of the wild allele (**Supplementary Fig. 8**). Similarly, *CHI1* (*Solyc05g010320*), encoding chalcone isomerase that catalyzes the committed step in the production of flavonoids, was consistently *cis*-regulated and favored the wild allele across tissues and developmental stages. In addition, *trans*-regulatory variation was detected specifically in pericarp tissue for genes such as *CHS2*, *F3’H*, *FLS,* and *Fd3GlcT*. Together with *cis* variants in flavonoid glycosylation genes, these findings expand the catalog of regulatory variants influencing polyphenolic diversity.

Domestication has markedly reduced bitter and toxic steroidal glycoalkaloids (SGAs) in modern tomatoes (Zhu et al., 2018). We detected predominant *trans*-regulation in early pathway genes that convert cholesterol to α-tomatine (**Fig. 6c**), with *S. pennellii* alleles showing elevated expression relative to *S. lycopersicum* alleles, whereas *S. neorickii* alleles were downregulated. By contrast, late-pathway genes (e.g., *GAME36*, *GAME5*), which catalyze the conversion of hydroxytomatine into non-bitter esculeoside A, showed strong *cis*-regulation, with cultivated alleles driving enhanced ripening-associated expression (**Supplementary Fig. 9**). These results suggest a domestication-linked regulatory shift: *trans*-regulation maintains α-tomatine biosynthesis in immature fruit, while *cis*-variants in cultivated alleles enhance its conversion into non-toxic, non-bitter metabolites during ripening.

Sugar composition represents a fundamental domestication trait that distinguishes cultivated tomatoes—which accumulate primarily fructose and glucose—from their wild relatives (e.g., *S. neorickii* and *S. pennellii*), which additionally accumulate sucrose (Husain et al., 2001). However, the *cis*– and *trans*-regulatory mechanisms governing sucrose accumulation and partitioning remain poorly understood. Our analysis of key sucrose metabolism genes revealed species-specific regulatory architectures underlying these metabolic differences. The vacuolar invertase gene *TIV1* (*Solyc03g083910*), which encodes an enzyme critical for sucrose hydrolysis (Klann et al., 1996), exhibited divergent regulatory pattern across species. It was predominantly *trans*-regulated in *S. pimpinellifolium* hybrids but shifted to near-constitutive *cis*-regulation in both *S. neorickii* and *S. pennellii* hybrids, with consistent upregulation of the cultivated allele across developmental stages (**Supplementary Fig. 10**). This regulatory divergence was complemented by allele expression differences in the invertase inhibitor gene *SlVIF* (*Solyc12g099190*), which inhibits sucrose hydrolysis during ripening (Qin et al., 2016). The wild *SlVIF* allele exhibited significantly higher expression than the cultivated allele in *S. neorickii* and *S. pennellii* hybrids during ripening–particularly at the ripe stage–thereby establishing a coordinated regulatory circuit that favors sucrose accumulation in green-fruited wild species (**Supplementary Fig. 10**). Together, these allele-specific expression patterns underscore the functional significance of *TIV1* and *SlVIF* in fine-tuning sucrose metabolism and establish that *cis*-regulatory divergence has been a primary mechanism underlying the optimization of sugar partitioning during tomato domestication and species evolution.

Volatile compounds are key determinants of tomato flavor and consumer preference. Our analysis revealed extensive *cis*-regulatory divergence in genes involved in the biosynthesis of volatiles derived from fatty acids, branched-chain amino acids (BCAAs), and nitrogen-containing precursors (**Supplementary Fig. 11a**). For fatty acid-derived volatiles, *LIP1* (*Solyc12g055730*) (Garbowicz et al., 2018) and *TomLoxC* (*Solyc01g006540*) (Shen et al., 2014) exhibited *cis* regulation during ripening. *LIP1*, encodes a lipase associated to the release of fatty acids from tryacylglycerols and was regulated by *cis* and *cis* + *trans* interactions in *S. neorickii* and *S. pennellii* hybrids during ripening favoring expression of the wild allele. *TomLoxC*, a lipoxygenase gene regulating fatty acid-derived volatile and apocarotenoid formation, exhibited a dynamic allelic expression shift during fruit development: in *S. pimpinellifolium* and *S. pennellii* hybrids the wild allele was preferentially expressed at early fruit stages, whereas the cultivated allele became dominant during ripening (**Supplementary Fig. 11**). This temporal shift aligns with previous findings that sequence variation in the *TomLoxC* promoter underlies transcriptional differences between wild and cultivated alleles (Gao et al., 2019; Li et al., 2023). In *S. neorickii* hybrids, however, wild alleles remained upregulated even during ripening, suggesting the involvement of distinct *cis*-regulatory elements in *S. neorickii*.

*SlBCAT1* (Solyc12g088220), which encodes a branched-chain aminotransferase involved in the biosynthesis of branched chain amino acid (BCAA)-derived volatiles (Maloney et al., 2010), showed near-constitutive cis regulation in *S. pennellii* hybrids with upregulation of the cultivated allele (**Supplementary Fig. 11a**). On the other hand, the *S. pimpinelifolium* allele of FLORAL4 (*Solyc04g063350*), which functions in the catabolism of BCAA and may also play a role in the production of phenolic-derived volatiles (Tikunov et al., 2020), was favored showing near-constitutive *cis*-regulation.

Additionally, several phenylpropanoid derived and nitrogen-containing volatile genes showed *cis*– and *cis +trans* regulatory divergence during ripening. *SlTNH1*, a gene involved in the ripening-related accumulation of nitrogenous volatiles (Liscombe et al., 2022) exhibited higher expression of the cultivated allele in *S. pennellii* hybrids. *SlMES1* (Solyc02g065240), encoding a methyl esterase involved in salicylic acid-derived volatile metabolism (Frick et al., 2023), showed upregulation of the *S. pimpinellifolium* and cultivated allele in *S. pimpinellifolium* and *S. neorickii* hybrids, respectively. However, The *S. neorickii* allele of *CTMOT1* (Solyc10g005060), a gene encoding a cathecol methyltransferase responsible for the synthesis of guaicol (Mageroy et al., 2012), was upregulated by *cis* effects (**Supplementary Fig. 11b**). Together, these results illustrate the complex and dynamic allele-specific regulatory landscape underlying volatile biosynthetic pathways in tomato hybrids, linking regulatory variation to metabolic and sensory diversity between cultivated tomatoes and their wild relatives.

## Discussion

Wild tomato species harbor remarkable morphological, metabolic, and genetic diversity, making them invaluable resources for tomato genetics and breeding (Meléndez-Martínez et al., 2010; Lin et al., 2014; Zhu et al., 2018; Li et al., 2023). The high-quality reference genomes of *S. pennellii* and *S. neorickii* generated in this study complement recently published wild tomato genomes (Wang et al., 2020; Li et al., 2023) and enable precise dissection of gene regulatory evolution. By integrating these high-quality genomes with high-resolution, tissue– and stage-resolved transcriptomes from interspecific crosses spanning diverse phylogenetic distances, we performed a comprehensive analysis of allele-specific expression in hybrids of cultivated tomato with three of its wild relatives. Recent studies in plants and animals have emphasized the role of tissue and cell type specificity in shaping the effects of regulatory variants (Andergassen et al., 2017; Ramasamy et al., 2025; Marand et al., 2025). However, comprehensive ASE analyses across tissues and developmental stages remain scarce in plants. Here, we included distinct fruit tissues at various stages of fruit development to enhance the power for capturing regulatory variation. Our results show a predominance of *cis*-regulated genes–particularly pronounced in hybrids involving the more distantly related wild species *S. neorickii* and *S. pennellii*–underscoring the central role of *cis* regulation in shaping phenotypic diversity during tomato evolution and domestication. Notably, a large number of genes exhibited *cis* regulation exclusively in a specific fruit tissue and /or developmental stage, highlighting the remarkable tissue specificity of the ASE process.

A prominent example of regulatory evolution is the extensive rewiring of ripening-associated metabolism required to convert bitter, chemically defended wild fruits into palatable, nutrient-rich cultivated tomatoes. This transition involves the coordinated downregulation of anti-nutritional steroidal glycoalkaloid (SGA) biosynthesis (Zhu et al., 2018; Liu et al., 2023a). Our analyses reveal a sophisticated dual regulatory pattern: *trans* regulation predominates in immature fruits to maintain defensive α-tomatine accumulation, whereas *cis*-regulatory variants in cultivated alleles promote the ripening-specific conversion of bitter tomatines into non-toxic esculeosides. This “defense early, edibility at maturity” switch exemplifies how ASE analyses can uncover the precise regulatory shifts that underlie tomato domestication.

Sugar metabolism is a hallmark domestication trait (Zhang et al., 2024). Our results reveal distinct *cis*-regulatory evolutionary trajectories in sucrose metabolism genes across tomato species. For example, we identified *cis*-regulated alleles shared by *S. neorickii* and *S. pennellii* in key metabolic genes, such as the vacuolar invertase gene *TIV1*, which catalyzes the hydrolysis of sucrose into fructose and glucose. Concurrently, the invertase inhibitor gene *SlVIF*, which binds to and suppresses *TIV1* activity, showed upregulation of the wild allele during ripening. This coordinated regulation reduces sucrose hydrolysis and thereby promotes sucrose accumulation in these green-fruited wild species. These allele-specific expression patterns underscore the functional significance of *TIV1* and *SlVIF* in fine-tuning sucrose metabolism. More broadly, they illustrate how *cis*-regulatory divergence contributes to the diversification of sugar profiles across tomato lineages, highlighting transcriptional reprogramming of sugar metabolism as a key feature of tomato evolution and domestication.

*Cis*-regulatory divergence is also evident in carotenoid biosynthesis, where cis-regulatory variants of *PSY1*, *LCYB*, and *LCYE* explain the evergreen fruit phenotype of *S. pennellii* and *S. neorickii*, in which lycopene synthesis is blocked and the pathway is diverted toward alternative carotenoids. Similarly, allelic differences in *PAL* and *CHI* may result in phenylalanine and flavonoid accumulation in *S. neorickii* and *S. pennellii*, consistent with QTLs identified in wild introgression lines (Brog et al., 2019; Szymański et al., 2020).

Fruit volatiles are critical determinants of consumer preference yet remain a major challenge in tomato breeding (Tieman et al., 2017; Klee and Tieman, 2018; Zhao et al., 2019). Extensive *cis*-regulatory divergence in genes involved in the biosynthesis of volatiles derived from fatty acids, branched-chain amino acids, and nitrogen-containing precursors provides a genetic blueprint for flavor diversification. Our results indicate that promoter SVs are likely a key driver of stable *cis*-regulatory divergence, particularly in more distantly related wild species. A similar mechanism has been reported for *TomLoxC*, in which a rare promoter allele greatly impacts tomato fruit flavor (Gao et al., 2019).

In conclusion, our study presents a high-spatiotemporal-resolution map of allele-specific expression underlying tomato fruit ripening. We establish *cis* regulation as a cornerstone of tomato evolution and domestication, shaping key quality traits such as sweetness, color, and flavor. By demonstrating how ASE analysis in interspecific hybrids can decode the evolutionary pattern of complex traits, this work advances our understanding of tomato domestication and identifies regulatory targets for future precision breeding aimed at improving tomato quality.

## Materials and Methods

### Genome sequencing, assembly, and pseudogenome construction

For *S. neorickii* LA2133 and *S. pennellii* LA0716, high-molecular-weight DNA was extracted from young fresh leaves using the cetyltrimethylammonium bromide (CTAB) method. PacBio SMRT libraries were constructed following the standard SMRTbell construction protocol (PacBio, USA) and sequenced on the PacBio Sequel II platform to generate HiFi reads. HiFi reads from each accession were assembled into contigs using hifiasm (Cheng et al., 2021) (v0.16.1-r375) with default parameters. The resulting contigs were then clustered into pseudochromosomes using RagTag (Alonge et al., 2022) (v2.1.0), with the *S. neorickii* ZY58 (Li et al., 2023) and *S. pennellii* LA0716 (Bolger et al., 2014b) genome assemblies serving as references for *S. neorickii* LA2133 and *S. pennellii* LA0716, respectively.

For *S. lycoperiscum* NC EBR-1 and TA209, genomic DNA was extracted from young fresh leaves using the DNeasy Plant Mini Kit (QIAGEN). Illumina libraries were constructed using the Illumina Genomic DNA Sample Preparation kit and sequenced on a NextSeq 500 platform. Raw reads were processed to trim adaptors and low-quality sequences using Trimmomatic (Bolger et al., 2014a) (v0.39) with parameters ‘TruSeq3-PE-2.fa:2:30:10:1:TRUE SLIDINGWINDOW:4:20 LEADING:3 TRAILING:3 MINLEN:40’. Cleaned reads were mapped to the ‘Heinz 1706’ reference genome (v4.0) (Hosmani et al., 2019) using BWA-MEM (Li, 2013) (v0.7.17) with default parameters. SNPs and small indels were then called using the Sentieon software package (https://www.sentieon.com/) and filtered using GATK (Mckenna et al., 2010) (v3.8) with parameters ‘QD < 2.0 || FS > 60.0 || MQ < 40.0 || SOR >3.0 || MQRankSum < –12.5 || ReadPosRankSum < –8.0’ for SNPs and ‘QD < 2.0 || FS > 200.0 || ReadPosRankSum < –20.0’ for small indels. The resulting high-quality variants were then patched and projected onto the ‘Heinz 1706’ genome to generate the final pseudogenomes of NC EBR-1 and TA209 using the g2gtools pipeline (https://github.com/churchill-lab/g2gtools).

### Repetitive sequence annotation and protein-coding gene prediction

Repetitive sequences in the *S. neorickii* LA2133 and *S. pennellii* LA0716 genome assemblies were annotated using EDTA (Ou et al., 2019) (v2.0.1) with parameters ‘--sensitive 1 –-anno 1’. Protein-coding genes were predicted from the repeat-masked genomes by integrating evidence from *ab initio*, homology-based, and transcriptome-based gene predictions. *Ab initio* gene predictions were performed using AUGUSTUS (Stanke et al., 2006) (v3.4.0), SNAP (Korf, 2004) (v2006-07-28) and GeneMark-ES (Lomsadze et al., 2005) (v4.69_lic). Homology-based evidence included protein sequences from tomato ‘Heinz 1706’ (v5.0) (Zhou et al., 2022), pepper Zhangshugang (Liu et al., 2023b), *S. pennellii* LA0716 (Bolger et al., 2014b) and other recently generated cultivated and wild tomato genomes (Li et al., 2023). Transcriptomic evidence was derived from RNA-seq data generated in this study and from NCBI (accession numbers SRR11044803, SRR3031964, SRR3119152, SRR3119156, SRR12714714, and SRR3031975). RNA-seq reads were processed using Trimmomatic (Bolger et al., 2014a) (v0.39) and *de novo* assembled using Trinity (Haas et al., 2013) (v2.14.0). Predictions from the above three approaches were integrated using Maker (Cantarel et al., 2008) (v3.01.04). To further improve gene prediction accuracy, gene models from ‘Heinz 1706’ (v5.0) (Zhou et al., 2022) and the recently generated cultivated and wild tomato genomes (Li et al., 2023) were mapped to all six parental genomes using Liftoff (Shumate and Salzberg, 2021) (v1.6.3) with parameters ‘-copies –sc 0.97 –exclude_partial –s 0.97 –a 0.95 –p 50’. Final gene models of the six parents were then generated using EVidenceModeler (Haas et al., 2008) (v1.1.1).

Protein-coding genes were functionally annotated by comparing their protein sequences against the NCBI non-redundant (nr), TrEMBL/SwissProt (Boeckmann et al., 2003), and InterPro (Mitchell et al., 2015) databases.

### Identification of unique gene-to-gene pairs between parental genomes

Unique gene-to-gene pairs between the two parents of each cross were identified by first mapping gene models from the cultivated parent to the wild parent genome using Liftoff (Shumate and Salzberg, 2021) (v1.6.3) with parameters ‘-copies –sc 0.97 –exclude_partial –s 0.97 –a 0.95 –p 50’. Genes from the wild parent that did not overlap mapped regions were then mapped back to the cultivated parent genome following the same procedure. Mapped gene models lacking valid open reading frames were still retained for further RNA-seq read mapping purpose. In addition, owing to the high synteny between the two parent genomes, only unique gene-to-gene pairs located on the same chromosomes were retained.

### Plant growth, RNA isolation, and sequencing

*Solanum lycopersicum* accessions M82, TA209, and NC EBR-1, *S. pimpinellifolium* LA2093, *S. pennellii* LA0716, and *S. neorickii* LA2133 were grown under standard greenhouse conditions with a 16 h/8 h light/dark photoperiod and day/night temperatures of 26 °C/18 °C. Reciprocal crosses between NC EBR-1 and LA2093 were performed using NC EBR-1 as either the pollen acceptor (NC EBR-1 × LA2093) or pollen donor (LA2093 × NC EBR-1). Crosses of M82 × LA0716 and TA209 × LA2133 were also performed, with wild parents used as pollen donors. Parents and F_1_ hybrids were grown under the same conditions described above, and hand pollinations were performed to ensure accurate dating of fruit age. For reciprocal crosses between NC EBR-1 and LA2093, fruits were collected at 7 dpa, mature green (MG), and ripe stages. For M82 × LA0716, fruits were collected at 21 dpa, MG, breaker (Br), and ripe stages. For TA209 × LA2133, fruits were collected at 7 dpa, MG, and ripe stages. At each stage, pericarp, placenta, and jelly tissues (when possible) were collected from 10 randomly selected individual plants per genotype. Total RNA was isolated from frozen tissues using the RNeasy Mini Kit (Qiagen). Strand-specific libraries were constructed following Zhong et al. (2011) and sequenced on an Illumina NextSeq 500 platform. Three biological replicates were conducted for each sample.

### Allele-specific expression quantification and regulatory mode classification

Raw RNA-seq reads were first processed using Trimmomatic (Bolger et al., 2014a) (v0.39) to remove adaptor and low-quality sequences with parameters ‘TruSeq3-PE-2.fa:2:30:10:1:TRUE SLIDINGWINDOW:4:20 LEADING:3 TRAILING:3 MINLEN:40’. Poly(A/T) tails were trimmed using PRINSEQ++ (Cantu et al., 2019) (v1.2.4) with parameters ‘-min_len 40 – trim_tail_left 10 –trim_tail_right 10’ and rRNA contaminations were removed by aligning reads to the SILVA rRNA database (Quast et al., 2013) (release 138) using HISAT2 (Kim et al., 2019) (v2.2.1). The resulting cleaned reads were mapped separately to the genomes of both parents using HISAT2. RNA-seq reads were then assigned to one of the two parents with higher mapping qualities or fewer mismatches. Reads with equal mapping qualities to both genomes were excluded from the downstream analyses.

Raw read counts for each unique gene-to-gene pair were calculated using featureCounts (Liao et al., 2014) (v2.0.3) with parameters ‘-p –s 2 –-countReadPairs –M –t gene’ and normalized to FPKM (fragments per kilobase of exon per million mapped fragments). Only genes with FPKM ≥ 1 in at least one sample of either parent or either parental allele in F_1_ hybrids were retained for regulatory mode classification. Two separate binomial exact tests were conducted using the DESeq2 R package (Love et al., 2014): one comparing the two parents (P1, cultivated; P2, wild) and the other comparing the parental alleles in F_1_ hybrids. Genes showing an adjusted *P*-value < 0.05 and fold change ≥ 2 or ≤ 0.5 were considered differentially expressed. To assign regulatory patterns (*cis*, *trans*, and their interactions), Fisher’s exact test (adjusted *P*-value < 0.05) was applied using expression values from the two parents and the two parental alleles in F_1_ hybrids. Based on these tests, genes were classified into five regulatory modes, following McManus et al. (2010) and Lemmon et al. (2014): (1) *cis*-regulated, if significant expression differences were observed between parents and between parental alleles in F_1_ hybrids but not in Fisher’s exact test; (2) *trans*-regulated, if significant expression differences were observed between parents and in Fisher’s exact test but not between parental alleles in F_1_ hybrids; (3) *cis* + *trans*-regulated if all three tests were significant and the favored allele showed increased expression in both parents and F_1_ hybrids; (4) *cis* × *trans*-regulated, if all three tests were significant but the favored allele showed increased expression in only one context (either parents or hybrids) but not both; and (5) compensatory if significant expression differences were observed between parental alleles in F_1_ hybrids and in Fisher’s exact test but not between the parents. All other patterns of significance or non-significance with no clear interpretation were named ambiguous.

The magnitude of *cis* effects was estimated by |log_2_ (hybrid P2 / hybrid P1)| and the magnitude of *trans* effects was calculated by subtracting *cis* effects from the total expression divergence between parental species: |log_2_ (parent P2 / parent P1) – |log_2_ (hybrid P2 / hybrid P1)|. The percentage of expression divergence contributed by *cis* effects was calculated as: %*cis* = *cis* magnitude / (*cis* magnitude + *trans* magnitude) × 100. The percentage of expression divergence contributed by *trans* effects was calculated as: %*trans* = *trans* magnitude / (*cis* magnitude + *trans* magnitude) × 100.

### Gene expression inheritance

Gene expression inheritance was determined by comparing expression levels in F1 hybrids with those in their parents. Genes whose expression in F_1_ hybrids was significantly lower than in P1 (cultivated parent) while significantly higher than in P2 (wild parent), or vice versa, were classified as additive. Genes whose expression in F_1_ hybrids differed significantly from P2 but not from P1 were classified as P1-dominant, whereas genes differing significantly from P1 but not from P2 were classified as P2-dominant. Genes whose expression in F_1_ hybrids was significantly higher than in both parents were classified as over-dominant, and those significantly lower than in both parents were classified as under-dominant.

### Variant identification among the six parents

Genomic variants, including SNPs, small indels, and SVs, were detected among the six parents using the Minigraph-Cactus pipeline (Hickey et al., 2024) with parameters ‘--permissiveContigFilter 0.1 –-maxLen 10000 –-clip 10000 –-vcf –-giraffe –-gfa –-gbz –-odgi’, using the M82 genome as the reference.

## Data availability

Raw RNA-seq reads have been deposited in the NCBI BioProject database under the accession number PRJNA1348326. Raw HiFi reads and genome assemblies have been deposited in the NCBI Bioproject database under the accession number PRJNA807529. Genome assemblies and annotations are also available at http://ted.bti.cornell.edu/ftp/tomato_genome/.

## Acknowledgements

This research was supported by grants from USDA National Institute of Food and Agriculture (2019-67013-29240) and the US National Science Foundation (IOS-1855585).

## Author contributions

C.C., Z.F. and J.J.G. designed and supervised the project. P.N., Y.X. and J.V. contributed to sample collection, DNA and RNA extraction and RNA-Seq library construction. Z.F. and C.C. coordinated genome and transcriptome sequencing. J.Z. performed genome assembly and annotation. J.Z. and P.N. contributed to transcriptome data processing and allele-specific expression analysis. J.Z., Z.F., and C.C. wrote the manuscript. Z.F. C.C. and P.N. revised the manuscript.

## Conflict of interest

The authors declare no conflict of interest.

## Notes

### Competing Interest Statement

The authors have declared no competing interest.

## References

1. Albert, E., Duboscq, R., Latreille, M., Santoni, S., Beukers, M., Bouchet, J.-P. P., Bitton, F., Gricourt, J., Poncet, C., Gautier, V., et al. (2018). Allele-specific expression and genetic determinants of transcriptomic variations in response to mild water deficit in tomato. Plant J. 96:635–650.

2. Alonge, M., Wang, X., Benoit, M., Soyk, S., Pereira, L., Zhang, L., Suresh, H., Ramakrishnan, S., Maumus, F., Ciren, D., et al. (2020). Major impacts of widespread structural variation on gene expression and crop improvement in tomato. Cell 182:145–161.e23.

3. Alonge, M., Lebeigle, L., Kirsche, M., Jenike, K., Ou, S., Aganezov, S., Wang, X., Lippman, Z. B., Schatz, M. C., and Soyk, S. (2022). Automated assembly scaffolding using RagTag elevates a new tomato system for high-throughput genome editing. Genome Biol. 23:258.

4. Alseekh, S., Ofner, I., Pleban, T., Tripodi, P., Di Dato, F., Cammareri, M., Mohammad, A., Grandillo, S., Fernie, A. R., and Zamir, D. (2013). Resolution by recombination: Breaking up *Solanum pennellii* introgressions. Trends Plant Sci. 18:536–538.

5. Andergassen, D., Dotter, C. P., Wenzel, D., Sigl, V., Bammer, P. C., Muckenhuber, M., Mayer, D., Kulinski, T. M., Theussl, H. C., Penninger, J. M., et al. (2017). Mapping the mouse Allelome reveals tissue-specific regulation of allelic expression. Elife 6:e25125.

6. Ashrafi, H., Kinkade, M. P., Merk, H. L., and Foolad, M. R. (2012). Identification of novel quantitative trait loci for increased lycopene content and other fruit quality traits in a tomato recombinant inbred line population. Mol. Breed. 30:549–567.

7. Bai, Y., and Lindhout, P. (2007). Domestication and breeding of tomatoes: What have we gained and what can we gain in the future? Ann. Bot. 100:1085–1094.

8. Bao, Y., Hu, G., Grover, C. E., Conover, J., Yuan, D., and Wendel, J. F. (2019). Unraveling *cis* and *trans* regulatory evolution during cotton domestication. Nat. Commun. 10:5399.

9. Bergougnoux, V. (2014). The history of tomato: From domestication to biopharming. Biotechnol. Adv. 32:170–189.

10. Boeckmann, B., Bairoch, A., Apweiler, R., Blatter, M. C., Estreicher, A., Gasteiger, E., Martin, M. J., Michoud, K., O’Donovan, C., Phan, I., et al. (2003). The SWISS-PROT protein knowledgebase and its supplement TrEMBL in 2003. Nucleic Acids Res. 31:365–370.

11. Bolger, A. M., Lohse, M., and Usadel, B. (2014a). Trimmomatic: A flexible trimmer for Illumina sequence data. Bioinformatics 30:2114–2120.

12. Bolger, A., Scossa, F., Bolger, M. E., Lanz, C., Maumus, F., Tohge, T., Quesneville, H., Alseekh, S., Sørensen, I., Lichtenstein, G., et al. (2014b). The genome of the stress-tolerant wild tomato species *Solanum pennellii*. Nat. Genet. 46:1034–1038.

13. Brog, Y. M., Osorio, S., Yichie, Y., Alseekh, S., Bensal, E., Kochevenko, A., Zamir, D., and Fernie, A. R. (2019). A *Solanum neorickii* introgression population providing a powerful complement to the extensively characterized *Solanum pennellii* population. Plant J. 97:391–403.

14. Cantarel, B. L., Korf, I., Robb, S. M. C., Parra, G., Ross, E., Moore, B., Holt, C., Alvarado, A. S., and Yandell, M. (2008). MAKER: An easy-to-use annotation pipeline designed for emerging model organism genomes. Genome Res. 18:188–196.

15. Cantu, V. A., Sadural, J., and Edwards, R. (2019). PRINSEQ++, a multi-threaded tool for fast and efficient quality control and preprocessing of sequencing datasets. PeerJ Prepr. 7:e27553v1.

16. Chakrabarti, M., Zhang, N., Sauvage, C., Muños, S., Blanca, J., Cañizares, J., Diez, M. J., Schneider, R., Mazourek, M., McClead, J., et al. (2013). A cytochrome P450 regulates a domestication trait in cultivated tomato. Proc. Natl. Acad. Sci. U. S. A. 110:17125–17130.

17. Chen, K. Y., Cong, B., Wing, R., Vrebalov, J., and Tanksley, S. D. (2007). Changes in regulation of a transcription factor lead to autogamy in cultivated tomatoes. Science 318:643–645.

18. Cheng, H., Concepcion, G. T., Feng, X., Zhang, H., and Li, H. (2021). Haplotype-resolved de novo assembly using phased assembly graphs with hifiasm. Nat. Methods 18:170–175.

19. Cong, B., Liu, J., and Tanksley, S. D. (2002). Natural alleles at a tomato fruit size quantitative trait locus differ by heterochronic regulatory mutations. Proc. Natl. Acad. Sci. U. S. A. 99:13606–13611.

20. Crowley, J. J., Zhabotynsky, V., Sun, W., Huang, S., Pakatci, I. K., Kim, Y., Wang, J. R., Morgan, A. P., Calaway, J. D., Aylor, D. L., et al. (2015). Analyses of allele-specific gene expression in highly divergent mouse crosses identifies pervasive allelic imbalance. Nat. Genet. 47:353–360.

21. Cubillos, F. A., Stegle, O., Grondin, C., Canut, M., Tisné, S., Gy, I., and Loudet, O. (2014). Extensive *cis*-regulatory variation robust to environmental perturbation in Arabidopsis. Plant Cell 26:4298–4310.

22. Doron-Faigenboim, A., Moy-Komemi, M., Petreikov, M., Eselson, Y., Sonawane, P., Cardenas, P., Fei, Z., Aharoni, A., and Schaffer, A. A. (2023). Transcriptomes of developing fruit of cultivated and wild tomato species. Mol. Hortic. 3:12.

23. Frick, E. M., Sapkota, M., Pereira, L., Wang, Y., Hermanns, A., Giovannoni, J. J., van der Knaap, E., Tieman, D. M., and Klee, H. J. (2023). A family of methyl esterases converts methyl salicylate to salicylic acid in ripening tomato fruit. Plant Physiol. 191:110–124.

24. Gao, L., Gonda, I., Sun, H., Ma, Q., Bao, K., Tieman, D. M., Burzynski-Chang, E. A., Fish, T. L., Stromberg, K. A., Sacks, G. L., et al. (2019). The tomato pan-genome uncovers new genes and a rare allele regulating fruit flavor. Nat. Genet. 51:1044–1051.

25. Garbowicz, K., Liu, Z., Alseekh, S., Tieman, D., Taylor, M., Kuhalskaya, A., Ofner, I., Zamir, D., Klee, H. J., Fernie, A. R., et al. (2018). Quantitative trait loci analysis identifies a prominent gene involved in the production of fatty acid-derived flavor volatiles in tomato. Mol. Plant 11:1147–1165.

26. Haas, B. J., Salzberg, S. L., Zhu, W., Pertea, M., Allen, J. E., Orvis, J., White, O., Buell, C. R., and Wortman, J. R. (2008). Automated eukaryotic gene structure annotation using EVidenceModeler and the Program to Assemble Spliced Alignments. Genome Biol. 9:R7.

27. Haas, B. J., Papanicolaou, A., Yassour, M., Grabherr, M., Blood, P. D., Bowden, J., Couger, M. B., Eccles, D., Li, B., Lieber, M., et al. (2013). De novo transcript sequence reconstruction from RNA-seq using the Trinity platform for reference generation and analysis. Nat. Protoc. 8:1494–1512.

28. Hickey, G., Monlong, J., Ebler, J., Novak, A. M., Eizenga, J. M., Gao, Y., Abel, H. J., Antonacci-Fulton, L. L., Asri, M., Baid, G., et al. (2024). Pangenome graph construction from genome alignments with Minigraph-Cactus. Nat. Biotechnol. 42:663–673.

29. Hosmani, P. S., Flores-Gonzalez, M., Geest, H. van de, Maumus, F., Bakker, L. V, Schijlen, E., Haarst, J. van, Cordewener, J., Sanchez-Perez, G., Peters, S., et al. (2019). An improved de novo assembly and annotation of the tomato reference genome using single-molecule sequencing, Hi-C proximity ligation and optical maps. bioRxiv 2012:767764.

30. Husain, S. E., James, C., Shields, R., and Foyer, C. H. (2001). Manipulation of fruit sugar composition but not content in *Lycopersicon esculentum* fruit by introgression of an acid invertase gene from *Lycopersicon pimpinellifolium*. New Phytol. 150:65–72.

31. Kim, D., Paggi, J. M., Park, C., Bennett, C., and Salzberg, S. L. (2019). Graph-based genome alignment and genotyping with HISAT2 and HISAT-genotype. Nat. Biotechnol. 37:907–915.

32. Klann, E. M., Hall, B., and Bennett, A. B. (1996). Antisense acid invertase (*TIV1*) gene alters soluble sugar composition and size in transgenic tomato fruit. Plant Physiol. 112:1321–1330.

33. Klee, H. J., and Tieman, D. M. (2018). The genetics of fruit flavour preferences. Nat. Rev. Genet. 19:347–356.

34. Koenig, D., Jiménez-Gómez, J. M., Kimura, S., Fulop, D., Chitwood, D. H., Headland, L. R., Kumar, R., Covington, M. F., Devisetty, U. K., Tat, A. V., et al. (2013). Comparative transcriptomics reveals patterns of selection in domesticated and wild tomato. Proc. Natl. Acad. Sci. U. S. A. 110: E2655–E2662.

35. Korf, I. (2004). Gene finding in novel genomes. BMC Bioinformatics 5:59.

36. Lemmon, Z. H., Bukowski, R., Sun, Q., and Doebley, J. F. (2014). The role of *cis* regulatory evolution in maize domestication. PLoS Genet. 10:e1004745.

37. Li, H. (2013). Aligning sequence reads, clone sequences and assembly contigs with BWA-MEM. arXiv 10.48550/arXiv.1303.3997.

38. Li, N., He, Q., Wang, J., Wang, B., Zhao, J., Huang, S., Yang, T., Tang, Y., Yang, S., Aisimutuola, P., et al. (2023). Super-pangenome analyses highlight genomic diversity and structural variation across wild and cultivated tomato species. Nat. Genet. 55:852–860.

39. Liao, Y., Smyth, G. K., and Shi, W. (2014). featureCounts: An efficient general purpose program for assigning sequence reads to genomic features. Bioinformatics 30:923–930.

40. Lin, T., Zhu, G., Zhang, J., Xu, X., Yu, Q., Zheng, Z., Zhang, Z., Lun, Y., Li, S., Wang, X., et al. (2014). Genomic analyses provide insights into the history of tomato breeding. Nat. Genet. 46:1220–1226.

41. Liscombe, D. K., Kamiyoshihara, Y., Ghironzi, J., Kempthorne, C. J., Hooton, K., Bulot, B., Kanellis, V., McNulty, J., Lam, N. B., Nadeau, L. F., et al. (2022). A flavin-dependent monooxygenase produces nitrogenous tomato aroma volatiles using cysteine as a nitrogen source. Proc. Natl. Acad. Sci. U. S. A. 119.

42. Liu, Y., Hu, H., Yang, R., Zhu, Z., and Cheng, K. (2023a). Current advances in the biosynthesis, metabolism, and transcriptional regulation of α-tomatine in tomato. Plants 12:3289.

43. Liu, F., Zhao, J., Sun, H., Xiong, C., Sun, X., Wang, X., Wang, Z., Jarret, R., Wang, J., Tang, B., et al. (2023b). Genomes of cultivated and wild Capsicum species provide insights into pepper domestication and population differentiation. Nat. Commun. 14:5487.

44. Lomsadze, A., Ter-Hovhannisyan, V., Chernoff, Y. O., and Borodovsky, M. (2005). Gene identification in novel eukaryotic genomes by self-training algorithm. Nucleic Acids Res. 33:6494–6506.

45. Love, M. I., Huber, W., and Anders, S. (2014). Moderated estimation of fold change and dispersion for RNA-seq data with DESeq2. Genome Biol. 15:550.

46. Mageroy, M. H., Tieman, D. M., Floystad, A., Taylor, M. G., and Klee, H. J. (2012). A *Solanum lycopersicum* catechol-O-methyltransferase involved in synthesis of the flavor molecule guaiacol. Plant J. 69:1043–1051.

47. Maloney, G. S., Kochevenko, A., Tieman, D. M., Tohge, T., Krieger, U., Zamir, D., Taylor, M. G., Fernie, A. R., and Klee, H. J. (2010). Characterization of the branched-chain amino acid aminotransferase enzyme family in tomato. Plant Physiol. 153:925–936.

48. Marand, A. P., Jiang, L., Gomez-Cano, F., Minow, M. A. A., Zhang, X., Mendieta, J. P., Luo, Z., Bang, S., Yan, H., Meyer, C., et al. (2025). The genetic architecture of cell type–specific *cis* regulation in maize. Science 388:eads6601.

49. Mckenna, A., Hanna, M., Banks, E., Sivachenko, A., Cibulskis, K., Kernytsky, A., Garimella, K., Altshuler, D., Gabriel, S., Daly, M., et al. (2010). The Genome Analysis Toolkit: A MapReduce framework for analyzing next-generation DNA sequencing data. Genome Res. 20:1297–1303.

50. McManus, C. J., Coolon, J. D., Duff, M. O., Eipper-Mains, J., Graveley, B. R., and Wittkopp, P. J. (2010). Regulatory divergence in *Drosophila* revealed by mRNA-seq. Genome Res. 20:816–825.

51. Meléndez-Martínez, A. J., Fraser, P. D., and Bramley, P. M. (2010). Accumulation of health promoting phytochemicals in wild relatives of tomato and their contribution to in vitro antioxidant activity. Phytochemistry 71:1104–1114.

52. Meyer, R. S., and Purugganan, M. D. (2013). Evolution of crop species: Genetics of domestication and diversification. Nat. Rev. Genet. 14:840–852.

53. Mitchell, A., Chang, H. Y., Daugherty, L., Fraser, M., Hunter, S., Lopez, R., McAnulla, C., McMenamin, C., Nuka, G., Pesseat, S., et al. (2015). The InterPro protein families database: The classification resource after 15 years. Nucleic Acids Res. 43:D213–D221.

54. Ou, S., Su, W., Liao, Y., Chougule, K., Agda, J. R. A., Hellinga, A. J., Lugo, C. S. B., Elliott, T. A., Ware, D., Peterson, T., et al. (2019). Benchmarking transposable element annotation methods for creation of a streamlined, comprehensive pipeline. Genome Biol. 20:275.

55. Qin, G., Zhu, Z., Wang, W., Cai, J., Chen, Y., Li, L., and Tian, S. (2016). A tomato vacuolar invertase inhibitor mediates sucrose metabolism and influences fruit ripening. Plant Physiol. 172:1596–1611.

56. Quast, C., Pruesse, E., Yilmaz, P., Gerken, J., Schweer, T., Yarza, P., Peplies, J., and Glöckner, F. O. (2013). The SILVA ribosomal RNA gene database project: Improved data processing and web-based tools. Nucleic Acids Res. 41:590–596.

57. Ramasamy, R., Raveendran, M., Harris, R. A., Le, H. D., Mure, L. S., Benegiamo, G., Dkhissi-Benyahya, O., Cooper, H., Rogers, J., and Panda, S. (2025). Genome-wide allele-specific expression in multi-tissue samples from healthy male baboons reveals the transcriptional complexity of mammals. Cell Genomics 5:100823.

58. Razifard, H., Ramos, A., Della Valle, A. L., Bodary, C., Goetz, E., Manser, E. J., Li, X., Zhang, L., Visa, S., Tieman, D., et al. (2020). Genomic evidence for complex domestication history of the cultivated tomato in Latin America. Mol. Biol. Evol. 37:1118–1132.

59. Rhie, A., Walenz, B. P., Koren, S., and Phillippy, A. M. (2020). Merqury: Reference-free quality, completeness, and phasing assessment for genome assemblies. Genome Biol. 21:245.

60. Rodríguez-Leal, D., Lemmon, Z. H., Man, J., Bartlett, M. E., and Lippman, Z. B. (2017). Engineering quantitative trait variation for crop improvement by genome editing. Cell 171:470–480.e8.

61. Sauvage, C., Rau, A., Aichholz, C., Chadoeuf, J., Sarah, G., Ruiz, M., Santoni, S., Causse, M., David, J., and Glémin, S. (2017). Domestication rewired gene expression and nucleotide diversity patterns in tomato. Plant J. 91:631–645.

62. Shao, L., Xing, F., Xu, C., Zhang, Q., Che, J., Wang, X., Song, J., Li, X., Xiao, J., Chen, L. L., et al. (2019). Patterns of genome-wide allele-specific expression in hybrid rice and the implications on the genetic basis of heterosis. Proc. Natl. Acad. Sci. U. S. A. 116:5653–5658.

63. Shen, J., Tieman, D., Jones, J. B., Taylor, M. G., Schmelz, E., Huffaker, A., Bies, D., Chen, K., and Klee, H. J. (2014). A 13-lipoxygenase, TomloxC, is essential for synthesis of C5 flavour volatiles in tomato. J. Exp. Bot. 65:419–428.

64. Shi, T., Gao, Z., Chen, J., and Van de Peer, Y. (2024). Dosage sensitivity shapes balanced expression and gene longevity of homoeologs after whole-genome duplications in angiosperms. Plant Cell 36:4323–4337.

65. Shumate, A., and Salzberg, S. L. (2021). Liftoff: Accurate mapping of gene annotations. Bioinformatics 37:1639–1643.

66. Signor, S. A., and Nuzhdin, S. V. (2018). The evolution of gene expression in *cis* and *trans*. Trends Genet. 34:532–544.

67. Simão, F. A., Waterhouse, R. M., Ioannidis, P., Kriventseva, E. V., and Zdobnov, E. M. (2015). BUSCO: Assessing genome assembly and annotation completeness with single-copy orthologs. Bioinformatics 31:3210–3212.

68. Soyk, S., Müller, N. A., Park, S. J., Schmalenbach, I., Jiang, K., Hayama, R., Zhang, L., Van Eck, J., Jiménez-Gómez, J. M., and Lippman, Z. B. (2017). Variation in the flowering gene *SELF PRUNING 5G* promotes day-neutrality and early yield in tomato. Nat. Genet. 49:162–168.

69. Stanke, M., Keller, O., Gunduz, I., Hayes, A., Waack, S., and Morgenstern, B. (2006). AUGUSTUS: *ab initio* prediction of alternative transcripts. Nucleic Acids Res. 34:435–439.

70. Swinnen, G., Goossens, A., and Pauwels, L. (2016). Lessons from domestication: Targeting *cis*-regulatory elements for crop improvement. Trends Plant Sci. 21:506–515.

71. Szymański, J., Bocobza, S., Panda, S., Sonawane, P., Cárdenas, P. D., Lashbrooke, J., Kamble, A., Shahaf, N., Meir, S., Bovy, A., et al. (2020). Analysis of wild tomato introgression lines elucidates the genetic basis of transcriptome and metabolome variation underlying fruit traits and pathogen response. Nat. Genet. 52:1111–1121.

72. Tieman, D., Zhu, G., Resende, M. F. R., Lin, T., Nguyen, C., Bies, D., Rambla, J. L., Beltran, K. S. O., Taylor, M., Zhang, B., et al. (2017). A chemical genetic roadmap to improved tomato flavor. Science 355:391–394.

73. Tikunov, Y. M., Roohanitaziani, R., Meijer-Dekens, F., Molthoff, J., Paulo, J., Finkers, R., Capel, I., Carvajal Moreno, F., Maliepaard, C., Nijenhuis-de Vries, M., et al. (2020). The genetic and functional analysis of flavor in commercial tomato: the FLORAL4 gene underlies a QTL for floral aroma volatiles in tomato fruit. Plant J. 103:1189–1204.

74. Tiroshauth, I., Reikhav, S., Levy, A. A., and Barkai, N. (2009). A yeast hybrid provides insight into the evolution of gene expression regulation. Science 324:659–662.

75. Wang, X., Gao, L., Jiao, C., Stravoravdis, S., Hosmani, P. S., Saha, S., Zhang, J., Mainiero, S., Strickler, S. R., Catala, C., et al. (2020). Genome of *Solanum pimpinellifolium* provides insights into structural variants during tomato breeding. Nat. Commun. 11:5817.

76. Wittkopp, P. J., and Kalay, G. (2012). *Cis*-regulatory elements: Molecular mechanisms and evolutionary processes underlying divergence. Nat. Rev. Genet. 13:59–69.

77. Wittkopp, P. J., Haerum, B. K., and Clark, A. G. (2004). Evolutionary changes in *cis* and *trans* gene regulation. Nature 430:85–88.

78. Xiao, H., Jiang, N., Schaffner, E., Stockinger, E. J., and Knaap, E. van der (2008). A retrotransposon-mediated gene duplication underlies morphological variation of tomato fruit. Science 319:1527–1530.

79. Zhang, J., Lyu, H., Chen, J., Cao, X., Du, R., Ma, L., Wang, N., Zhu, Z., Rao, J., Wang, J., et al. (2024). Releasing a sugar brake generates sweeter tomato without yield penalty. Nature 635:647–656.

80. Zhao, J., Sauvage, C., Zhao, J., Bitton, F., Bauchet, G., Liu, D., Huang, S., Tieman, D. M., Klee, H. J., and Causse, M. (2019). Meta-analysis of genome-wide association studies provides insights into genetic control of tomato flavor. Nat. Commun. 10:1534.

81. Zhong, S., Joung, J. G., Zheng, Y., Chen, Y. R., Liu, B., Shao, Y., Xiang, J. Z., Fei, Z., and Giovannoni, J. J. (2011). High-throughput Illumina strand-specific RNA sequencing library preparation. Cold Spring Harb. Protoc. 6:940–949.

82. Zhou, Y., Zhang, Z., Bao, Z., Li, H., Lyu, Y., Zan, Y., Wu, Y., Cheng, L., Fang, Y., Wu, K., et al. (2022). Graph pangenome captures missing heritability and empowers tomato breeding. Nature 606:527–534.

83. Zhu, G., Wang, S., Huang, Z., Zhang, S., Liao, Q., Zhang, C., Lin, T., Qin, M., Peng, M., Yang, C., et al. (2018). Rewiring of the fruit metabolome in tomato breeding. Cell 172:249–261.e12.

