## Supplementary Figures for "Allele-specific expression reveals complex *cis*- and *trans*-regulatory divergence underlying fruit phenotypic differences between cultivated and wild tomato species"

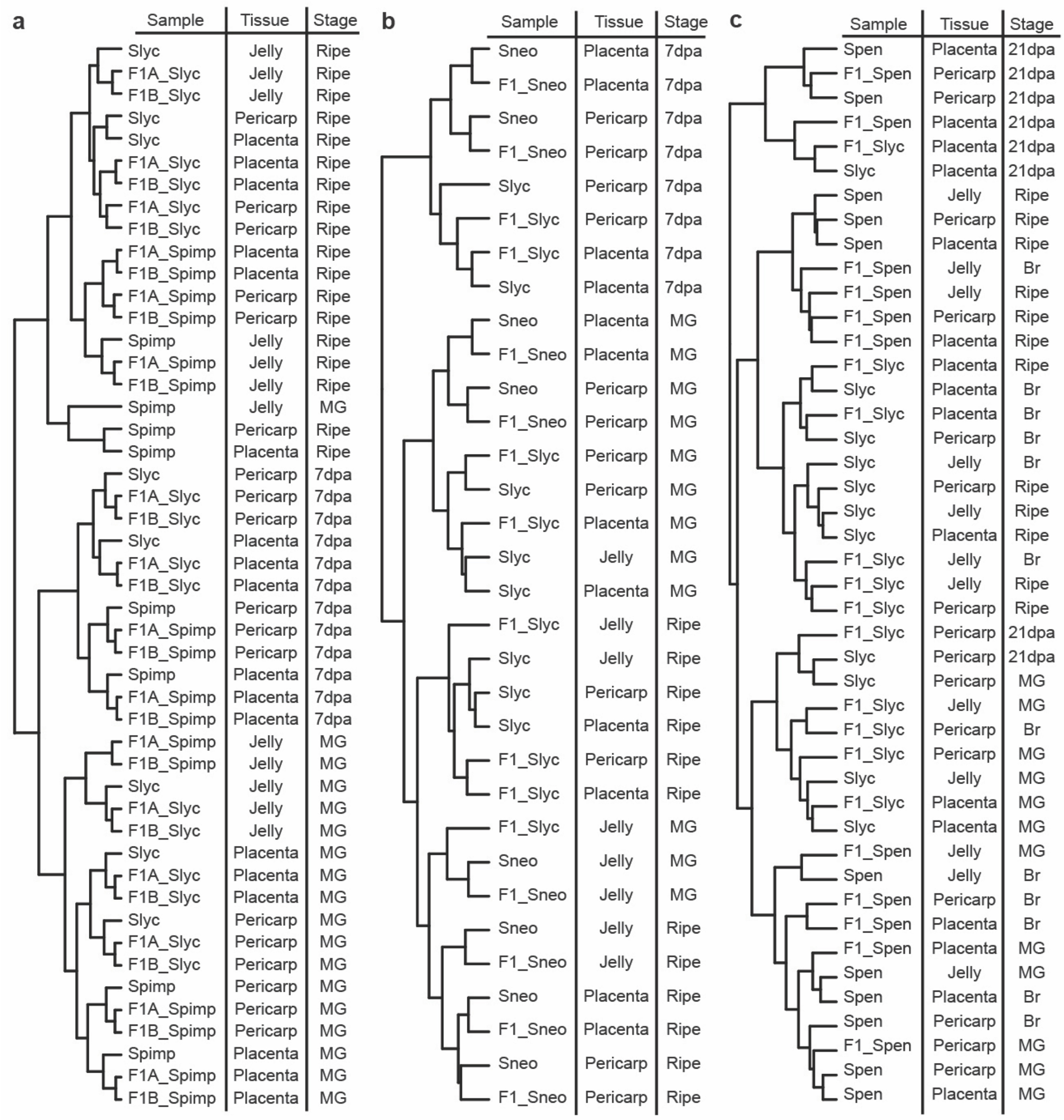


**Supplementary Fig. 1 Sample correlation analysis of allele-specific gene expression.** **a**, Clustering of samples from *S. lycopersicum* NC EBR-1 × *S. pimpinellifolium* LA2093 reciprocal hybrids and their parental lines. F1A, $F_1$ from NC EBR-1 (♀) × LA2093 (♂); F1B, $F_1$ from LA2093 (♀) × NC EBR-1 (♂). **b**, Clustering of samples from *S. lycopersicum* TA209 × *S. neorickii* LA2133 and their parental lines. **c**, Clustering of samples from *S. lycopersicum* M82 × *S. pennellii* LA0716 and their parental lines. Slyc, *S. lycopersicum*; Spimp, *S. pimpinellifolium*; Sneo, *S. neorickii*; Spen, *S. pennellii*. dpa, days post anthesis; MG, mature green; Br, breaker.

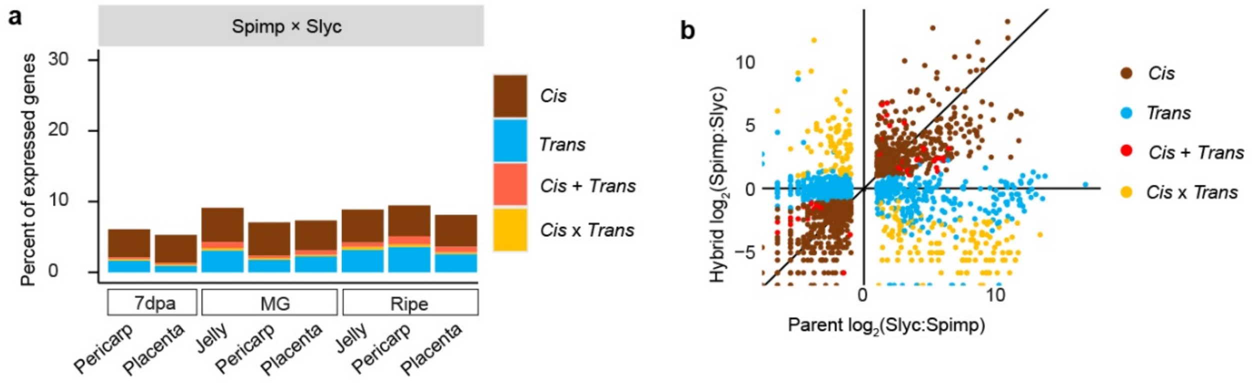


**Supplementary Fig. 2 Allele-specific expression in *S. pimpinellifolium* LA2093 (♀) × *S. lycopersicum* NC EBR-1 (♂) hybrids**. **a**, Percentage of genes in different regulatory categories across fruit tissues and developmental stages. **b**, Scatter plots showing the relative expression of the cultivated and wild alleles in parents versus the relative expression of the corresponding alleles in the $F_1$ hybrid for the pericarp at the ripe stage. dpa, days post anthesis; MG, mature green; Br, breaker. Slyc, *S. lycopersicum*; Spimp, *S. pimpinellifolium*.

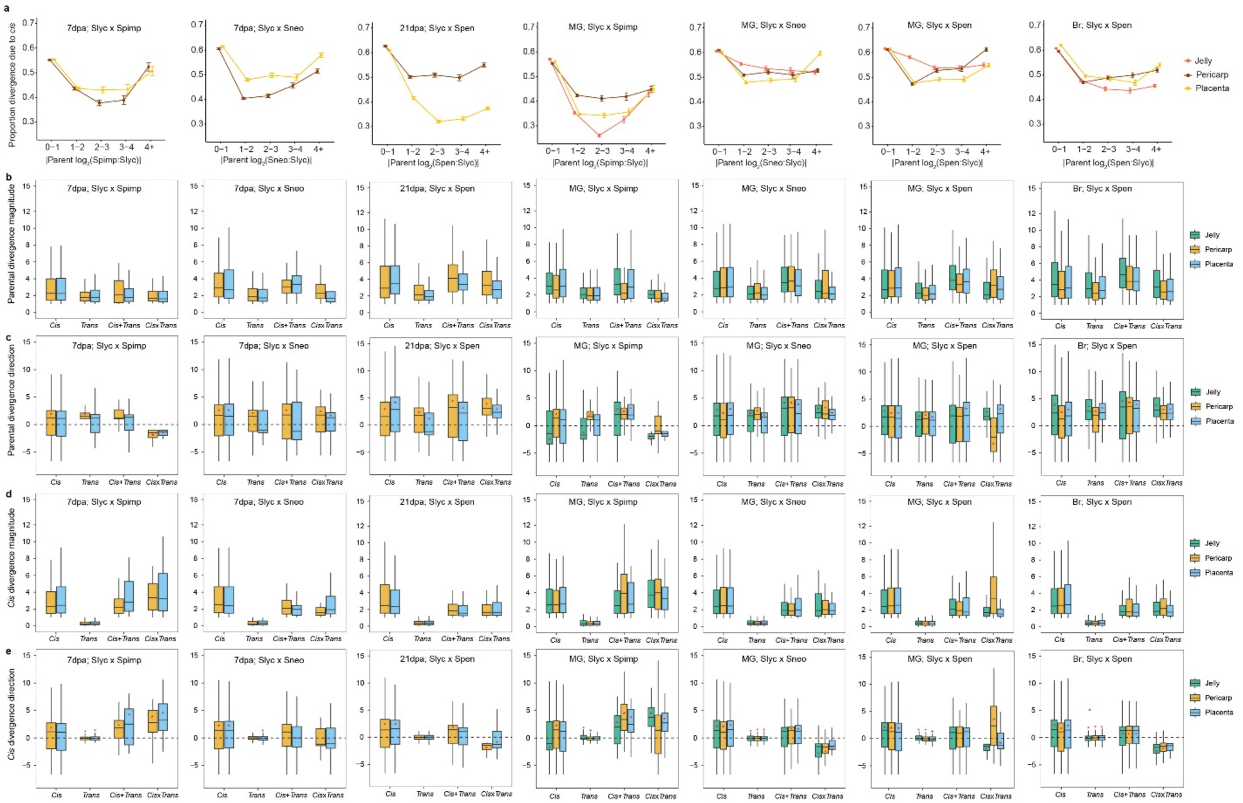


**Supplementary Fig. 3 Expression divergence attributable to *cis* regulation at 7 dpa, 21 dpa, MG, and Br stages. a**, Proportion of parental expression divergence explained by *cis*-regulatory effects at 7 dpa in *S. lycopersicum* × *S. pimpinellifolium* and *S. lycopersicum* × *S. neorickii*; at 21 dpa in *S. lycopersicum* × *S. pennellii*; at MG in *S. lycopersicum* × *S. pimpinellifolium*, *S. lycopersicum* × *S. neorickii*, and *S. lycopersicum* × *S. pennellii*; and at Br in *S. lycopersicum* × *S. pennellii* (from left to right). Genes were binned according to parental expression divergence, as shown on the x-axis. **b-e**, Parental divergence magnitude ($|\log_2(P2/P1)|$) (**b**), parental divergence direction ($\log_2(P2/P1)$) (**c**), *cis* divergence magnitude ($|\log_2(P2/P1)|$ in $F_1$) (**d**), and *cis* divergence direction ($\log_2(P2/P1)$ in $F_1$) (**e**) for genes under *cis*, *trans*, *cis* + *trans*, or *cis* × *trans* regulation. Data are shown for the same developmental stages and species combinations as in panel **a** (from left to right). Significant deviations from zero, indicated by red stars (*), were inferred using the Wilcoxon test ($P < 0.05$).

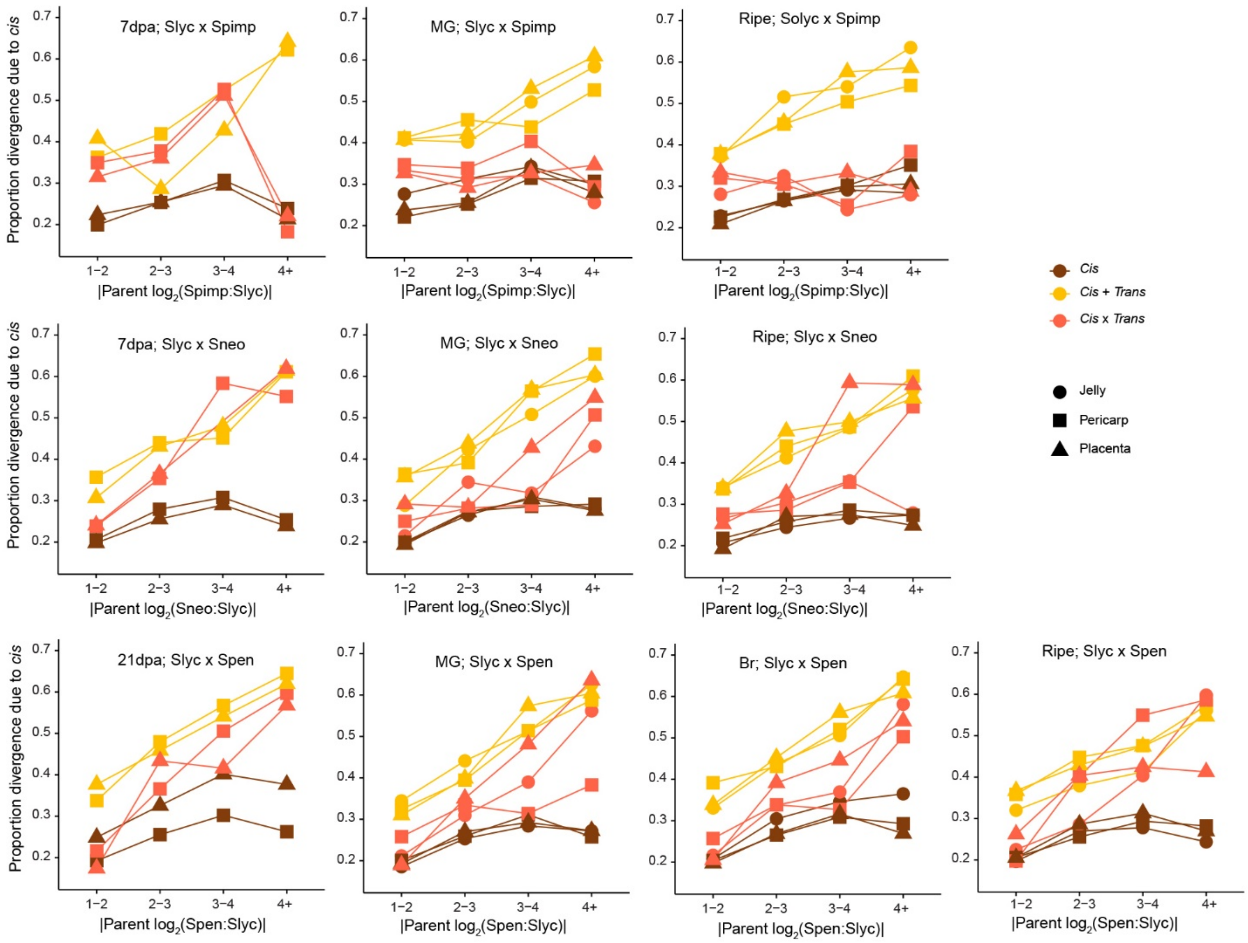


**Supplementary Fig. 4** Proportion of parental expression divergence explained by *cis*-regulatory effects at different developing stages/tissues for *cis*, *cis* + *trans* and *cis* × *trans* genes. Slyc × Spimp, *S. lycopersicum* × *S. pimpinellifolium*; Slyc × Sneo, *S. lycopersicum* × *S. neorickii*; Slyc × Spen, *S. lycopersicum* × *S. pennellii*. MG, mature green; Br, breaker. Genes were binned according to parental expression divergence, as shown on the x-axis.

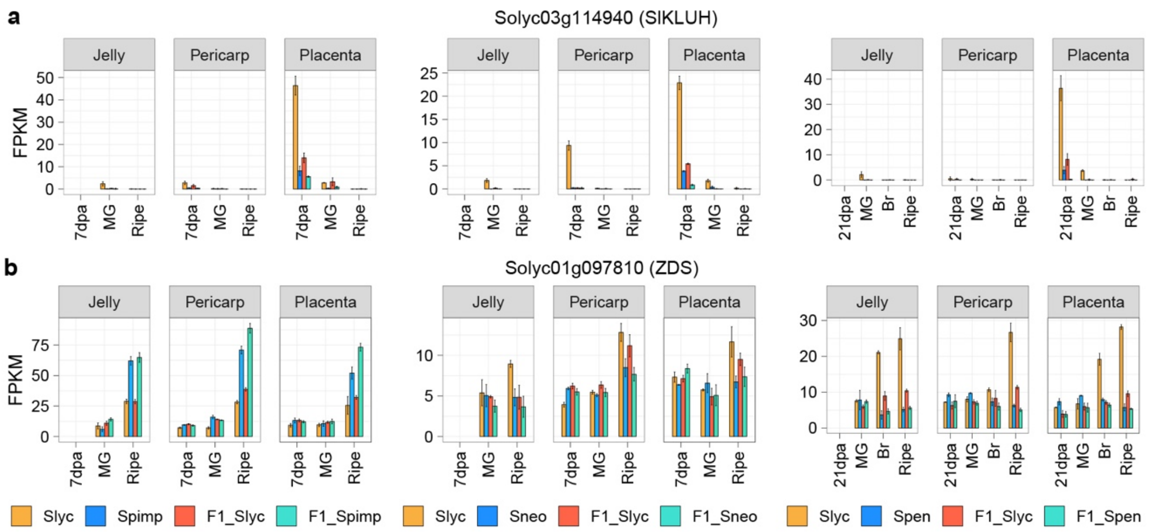


**Supplementary Fig. 5 Expression of tissue-specific *cis*-regulated genes *SlKLUH* (a) and *ZDS* (b)**. Although cultivated and wild alleles of *ZDS* showed differential expression in jelly and placenta, these differences did not meet the statistical criteria for *cis* regulation. Slyc, *S. lycopersicum*; Spimp, *S. pimpinellifolium*; Sneo, *S. neorickii*; Spen, *S. pennellii*. dpa, days post anthesis; MG, mature green; Br, breaker.

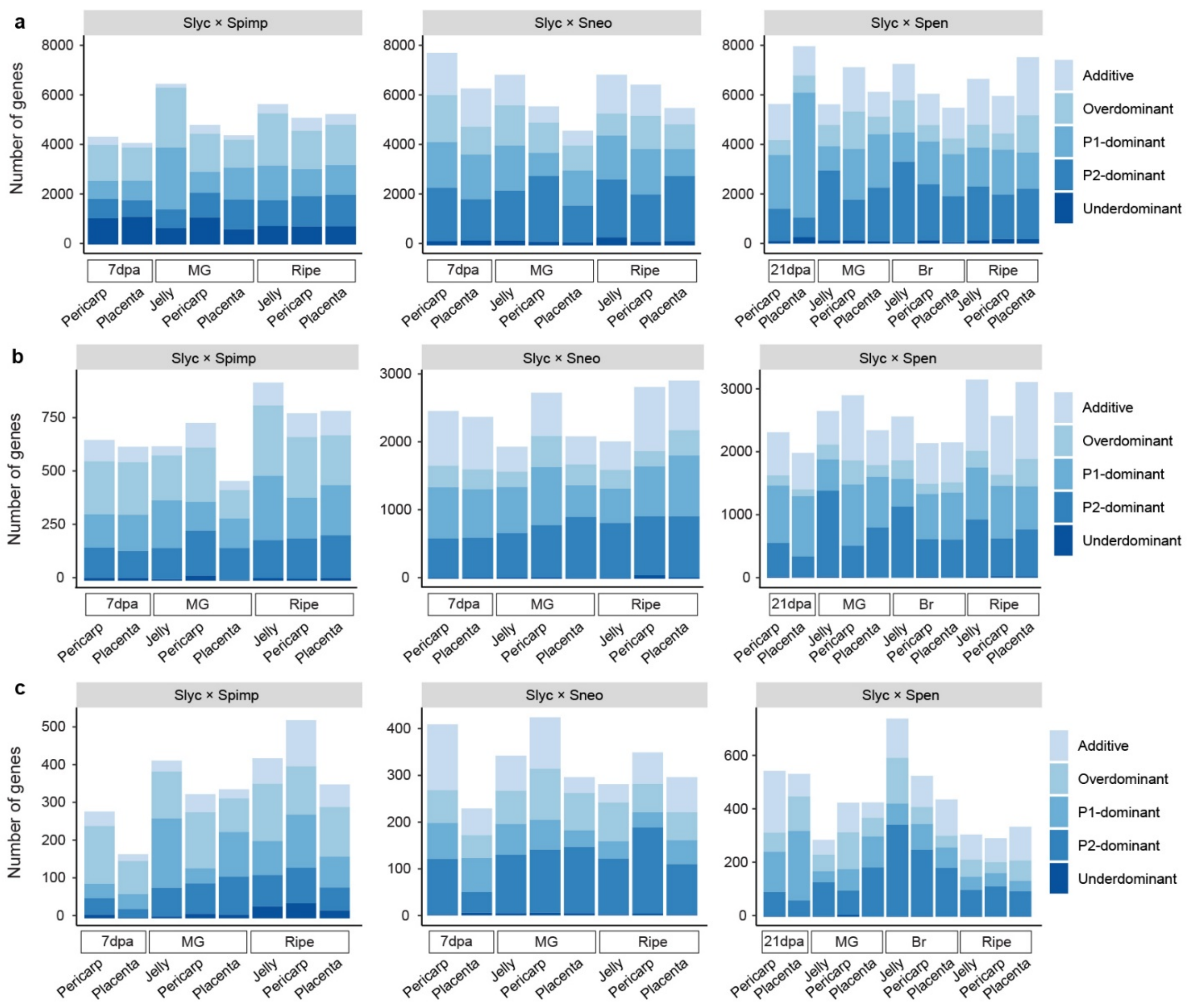


**Supplementary Fig. 6 Gene expression inheritance in tomato hybrids. a**, Number of genes in each inheritance category across fruit tissues and developmental stages. P1, cultivated tomato parent; P2, wild tomato parent. **b,c**, Distribution of inheritance modes among *cis*- (**b**) and *trans*-regulated genes (**c**). Within each inheritance category, bars represents individual stages or tissues, shown in the same order as in panel **a**. dpa, days post anthesis; MG, mature green; Br, breaker. Slyc, *S. lycopersicum*; Spimp, *S. pimpinellifolium.*

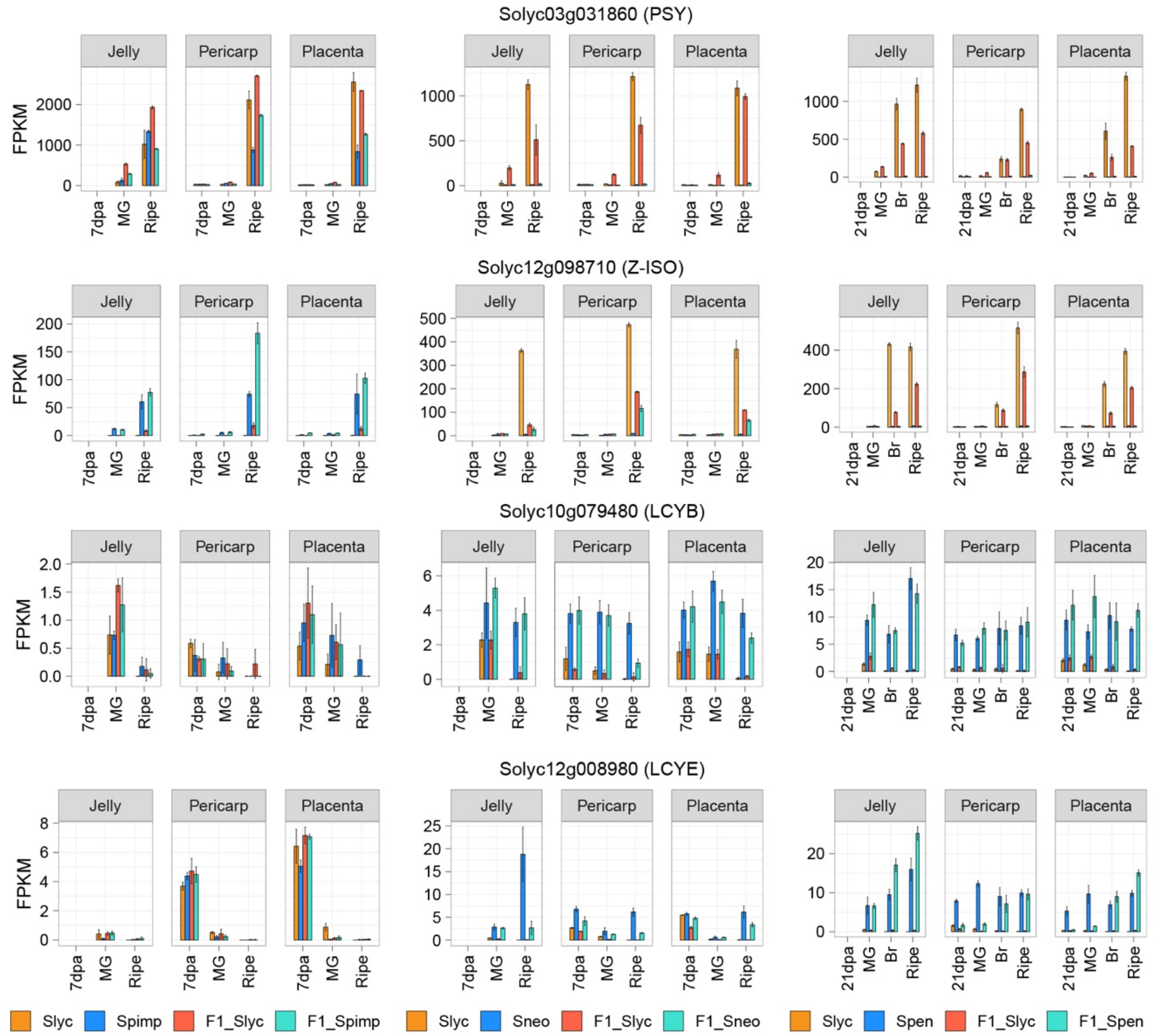


**Supplementary Fig. 7 Allele-specific expression of genes involved in carotenoid biosynthesis**. Slyc, *S. lycopersicum*; Spimp, *S. pimpinellifolium*; Sneo, *S. neorickii*; Spen, *S. pennellii*. dpa, days post anthesis; MG, mature green; Br, breaker. PSY1, phytoene synthase 1; Z-ISO, ζ-carotene isomerase; LCYB, lycopene β-cyclase; LCYE, lycopene ε-cyclase.

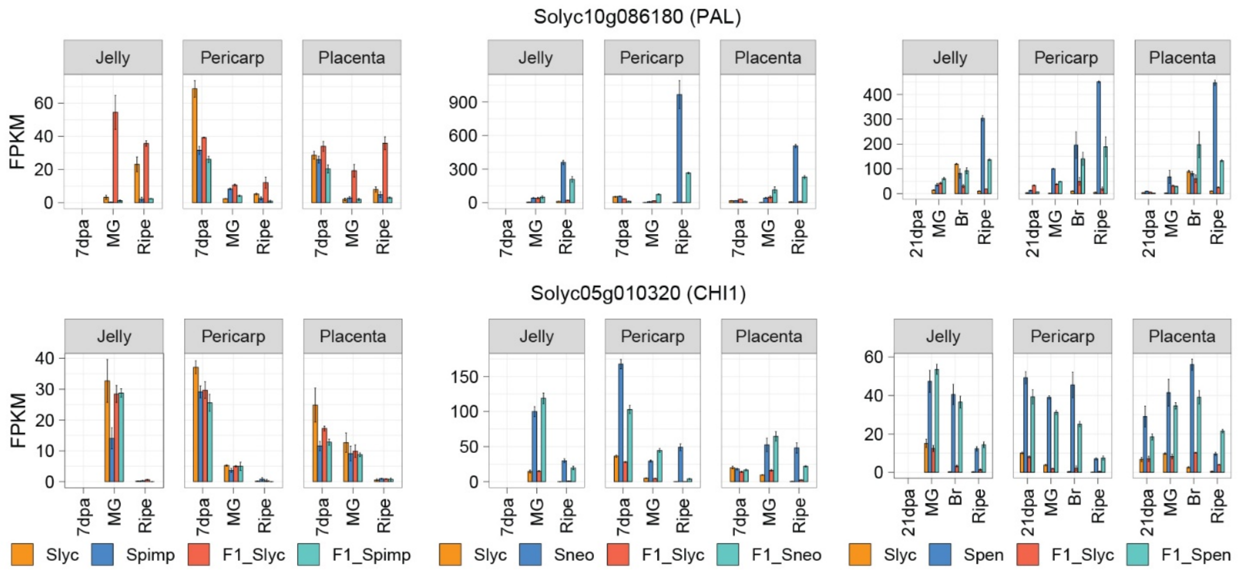


**Supplementary Fig. 8 Allele-specific expression of genes involved in phenylpropanoid and flavonoid biosynthesis.** Slyc, *S. lycopersicum*; Spimp, *S. pimpinellifolium*; Sneo, *S. neorickii*; Spen, *S. pennellii*. dpa, days post anthesis; MG, mature green; Br, breaker. PAL, phenylalanine ammonia-lyase; CHI, chalcone isomerase.

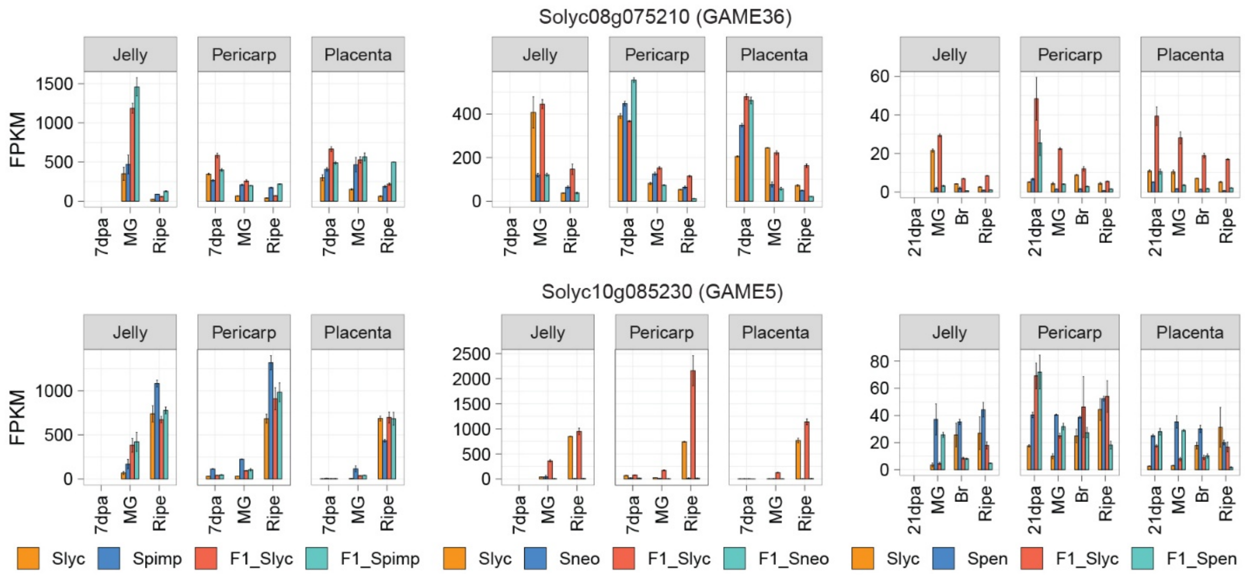


**Supplementary Fig. 9 Allele-specific expression of genes involved in steroidal glycoalkaloid metabolism.** Slyc, *S. lycopersicum*; Spimp, *S. pimpinellifolium*; Sneo, *S. neorickii*; Spen, *S. pennellii*. dpa, days post anthesis; MG, mature green; Br, breaker. GAME, glycoalkaloid metabolism.

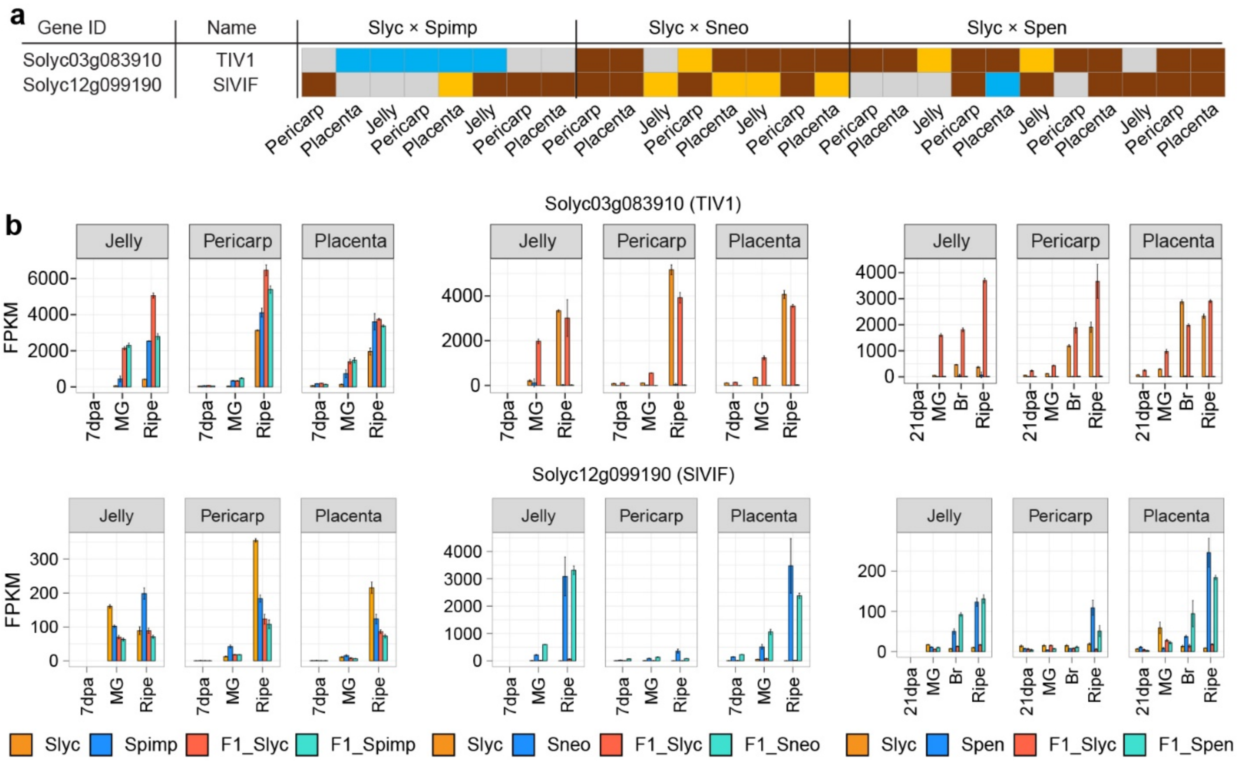


**Supplementary Fig. 10 Regulatory and expression patterns of genes involved in sugar metabolisum in tomato hybrids**. **a,b**, Regulatory patterns (**a**) and expression profiles (**b**) of genes involved in sugar metabolism. *TIV1*, acid invertase; *SlVIF*, vacuolar invertase. Slyc, *S. lycopersicum*; Spimp, *S. pimpinellifolium*; Sneo, *S. neorickii*; Spen, *S. pennellii*. dpa, days post anthesis; MG, mature green; Br, breaker.

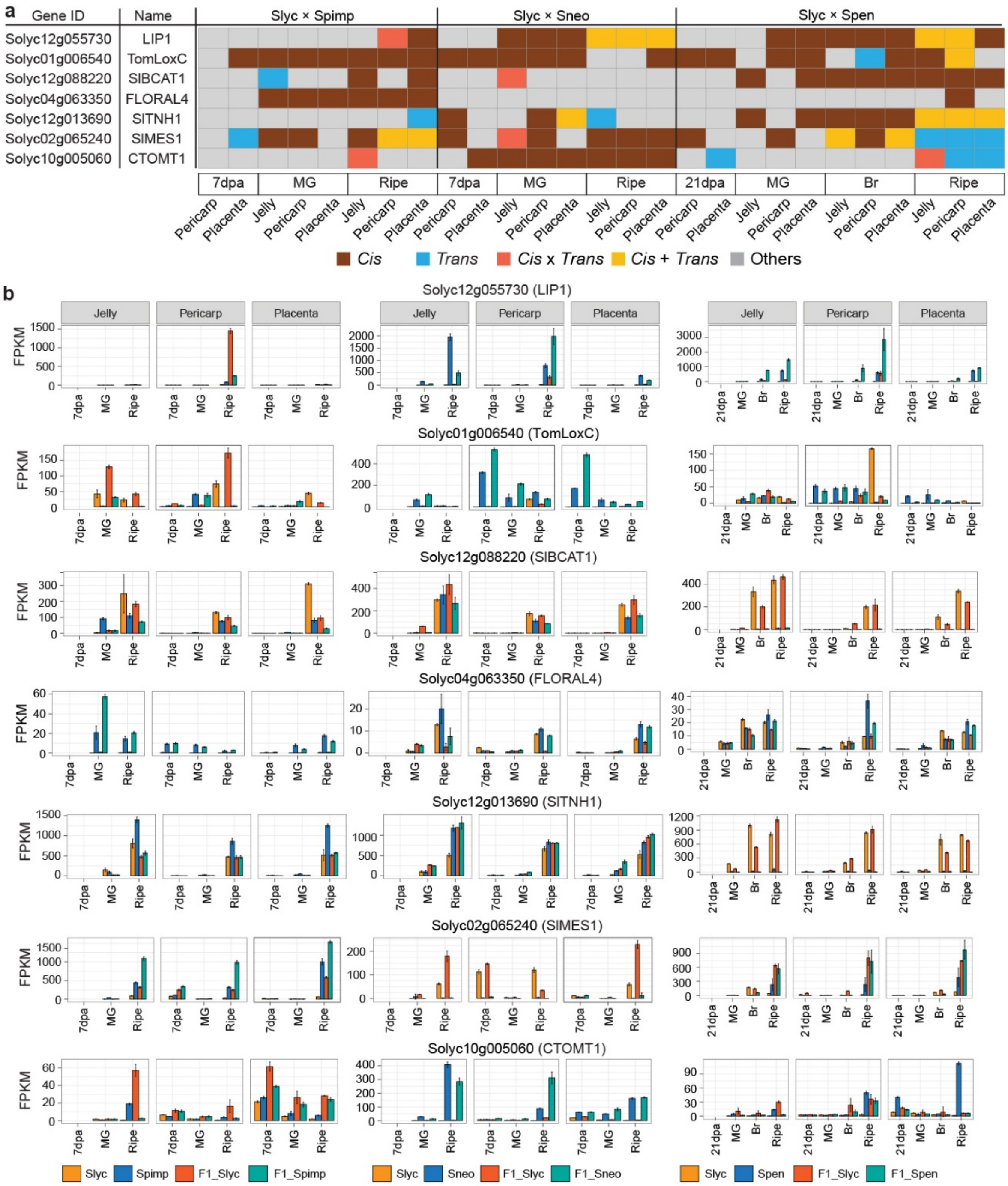


**Supplementary Fig. 11 Regulatory patterns and expression of genes involved in volatiles in tomato hybrids**. **a**, Regulatory patterns (**a**) and expression profiles (**b**) of genes involved in volatile biosynthesis. LIP1, class III lipase 1; TomLoxC, tomato 13-lipoxygenase C; SlBCAT1, tomato branched-chain amino acid aminotransferase 1; FLORAL4, 2-oxoisovalerate dehydrogenase; SlTNH1, flavin-dependent monooxygenase; SlMES1, salicylic acid methyl esterase 1; CTOMT1, catechol-O-methyltransferase. Slyc, *S. lycopersicum*; Spimp, *S. pimpinellifolium*; Sneo, *S. neorickii*; Spen, *S. pennellii*. dpa, days post anthesis; MG, mature green; Br, breaker.
